# Molecular mechanism for the recognition of sequence-divergent CIF peptides by the plant receptor kinases GSO1/SGN3 and GSO2

**DOI:** 10.1101/692228

**Authors:** Satohiro Okuda, Satoshi Fujita, Andrea Moretti, Ulrich Hohmann, Verónica G. Doblas, Yan Ma, Alexandre Pfister, Benjamin Brandt, Niko Geldner, Michael Hothorn

**Author notes:** Institut Jean-Pierre Bourgin, INRA, AgroParisTech, CNRS, Université Paris-Saclay, 78000 Versailles, France. National Institute of Genetics, Mishima, Shizuoka 411-8540, Japan (Satoshi Fujita). Institute of Molecular Biotechnology of the Austrian Academy of Sciences (IMBA) & Research Institute of Molecular Pathology (IMP), Vienna Biocenter (VBC), 1030 Vienna, Austria. Department of Plant and Microbial Biology, University of Zurich, 8008 Zurich, Switzerland (Benjamin Brandt). These authors contributed equally to this work.

## Abstract

Plants use leucine-rich repeat receptor kinases (LRR-RKs) to sense sequence diverse peptide hormones at the cell surface. A 3.0 Å crystal structure of the LRR-RK GSO1/SGN3 regulating Casparian strip formation in the endodermis reveals a large spiral-shaped ectodomain. The domain provides a binding platform for 21 amino-acid CIF peptide ligands, which are tyrosine sulfated by the tyrosylprotein sulfotransferase TPST/SGN2. GSO1/SGN3 harbors a binding pocket for sulfotyrosine and makes extended backbone interactions with CIF2. Quantitative biochemical comparisons reveal that GSO1/SGN3 – CIF2 represents one of the strongest receptor-ligand pairs known in plants. Multiple missense mutations are required to block CIF2 binding *in vitro*, and GSO1/SGN3 function *in vivo*. Using structure-guided sequence analysis we uncover novel CIF peptides conserved among higher plants. Quantitative binding assays with known and novel CIFs suggest that the homologous LRR-RKs GSO1/SGN3 and GSO2 have evolved unique peptide binding properties to control different developmental processes. A quantitative biochemical interaction screen, a CIF peptide antagonist and genetic analyses together implicate SERK LRR-RKs as essential co-receptor kinases required for GSO1/SGN3 and GSO2 receptor activation. 0ur work provides a mechanistic framework for the recognition of sequence-divergent peptide hormones in plants.

**Significance Statement:** Two sequence-related plant membrane receptor kinases and their shape-complementary co-receptors are shown to selectively sense members of a small family of secreted peptide hormones to control formation of an important diffusion barrier in the plant root.

## Introduction

Plant membrane receptor kinases with leucine-rich repeat ectodomains (LRR-RKs) form the first layer of the plant immune system and are key regulators of plant growth and development (1). LRR-RKs have evolved to sense small molecule, peptide and protein ligands, with small linear peptides representing a large class of sequence-diverse signaling molecules in plants (1, 2). These linear peptides are processed from larger pre-proteins and subsequently post-translationally modified (3). The size of the final, bioactive peptide hormone ranges from five (phytosulfokine, PSK) (2) to ~ 21-23 amino-acids (PEP1; CASPARIAN STRIP INTEGRITY FACTORS, CIF1/2) (4–6). Post-translational peptide modifications include proline hydroxylation, hydroxyproline arabinosylation, and tyrosine sulfation (sTyr) (2), and these modifications may allow for specific ligand recognition by the cognate LRR-RK (7–9). The disulfated PSK peptide binds to a pocket that is formed by the LRR domain of the receptor PSKR and a small ‘island domain’ (9). PSK binding stabilizes the island domain and enables PSKR to interact with a SERK co-receptor kinase, which is shared between many LRR-RK signaling pathways (9, 1). Unsulfated PSK variants bound the receptor with ~25fold reduced affinity (9). Subsequently, other tyrosine sulfated peptides were discovered in plants, including the ROOT MERISTEM GROWTH FACTORS (RGFs), 13 amino-acid peptides containing an N-terminal Asp-Tyr (DY) motif (10), which is recognized by the sole tyrosylprotein sulfotransferase TPST in Arabidopsis (11). RGFs are sensed by a class of SERK-dependent LRR-RKs termed RGFRs (12, 13). RGFs bind the LRR ectodomain of RGFRs with dissociation constants in the high nanomolar range (13). Non-sulfated variants of the linear peptides showed a ~200fold reduction in binding affinity (13). The N-terminal sTyr in RGFs maps to a hydrophobic pocket located at the inner face of the LRR solenoid in RGF-RGFR complex structures, with the peptide adopting an extended conformation (13). A His-Asn diad forms the C-terminus of RGFs and many other plant peptide hormones, such as IDA/IDLs involved in organ abscission and CLE peptides controlling plant stem cell maintenance (7, 1). The C-terminal His/Asn motif has been shown to be specifically recognized by two arginines (the RxR motif) located at the inner surface of the LRR cores of different peptide sensing LRR-RKs (7, 13–16).

The LRR-RKs GASSHO1/SCHENGEN 3 (GSO1/SGN3) and GASSHO2 (GSO2) carry a conserved RxR motif and were initially shown to be redundantly required for embryonic development (17, 18). Subsequently, a non-redundant role for GSO1/SGN3 was identified through a genetic screen for Casparian strip formation, an endodermal barrier allowing for selective nutrient uptake in the root (19, 20). The presence of the RxR motif suggested that GSO1/SGN3 and GSO2 may bind peptide ligands *in planta*, but the identify of these peptides remained unknown. The discovery that *tpst*/*sgn2* loss-of-function mutants display Casparian strip phenotypes similar to *sgn3* resulted in the identification of two 21 amino-acid long, tyrosine sulfated peptides CIF1/2 as ligands for GSO1/SGN3 (6). A complementary biochemical interaction screen for CIF1/2 receptors identified GSO1/SGN3 and GSO2 as *bona fide* receptors for these peptide hormones (5). Here we report the crystal structure of the GSO1/SGN3 ectodomain in complex with CIF2 and dissect its mode of ligand binding. We define novel CIF peptides differentially sensed by GSO1/SGN3 and GSO2 and report that GSO1 and GSO2 require SERK co-receptor kinases for receptor activation.

## Results

The interaction between the GSO1/SGN3 ectodomain and synthetic CIF1/2 peptides has been previously characterized in quantitative isothermal titration calorimetry (ITC) steady-state binding assays, yielding dissociation constants (K_*d*_’s) ranging from ~2 to 50 nM, but with varying binding stoichiometries (6). We performed grating coupled interferometry (GCI) kinetic binding assays (21) and found that GSO1/SGN3 binds the CIF1 and CIF2 peptides with K_*d*_’s of ~5 and ~1 nM, respectively (Fig. 1), in agreement with the earlier report (6). Next, we compared the binding kinetics of GSO1/SGN3 – CIF1/CIF2 to other, known receptor – peptide ligand pairs from Arabidopsis: The 23 amino-acid PEP1 and PEP2 danger signal peptides bind the LRR-RK PEPR1 with drastically different binding affinities of 90 nM and 18 μM, respectively (Fig. 1). The hydroxyprolinated CLE9 peptide (12 amino-acids) binds the ectodomain of the LRR-RK BAM1 with a K_*d*_ of ~1 nM, similar to GSO1/SGN3 – CIF2 (Fig. 1), and in agreement with a previously reported ITC experiment (22). The well-characterized immune elicitor peptide flg22 binds the isolated FLS2 ectodomain with a dissociation constant of 1.5 μM (Fig. 1). Together, our comparison reveals that plant LRR-RKs can sense peptide ligands with drastically different binding affinities and kinetics, with the GSO1/SGN3 – CIF1/2 interaction ranking among the strongest receptor – ligand pairs.

**Fig. 1.**
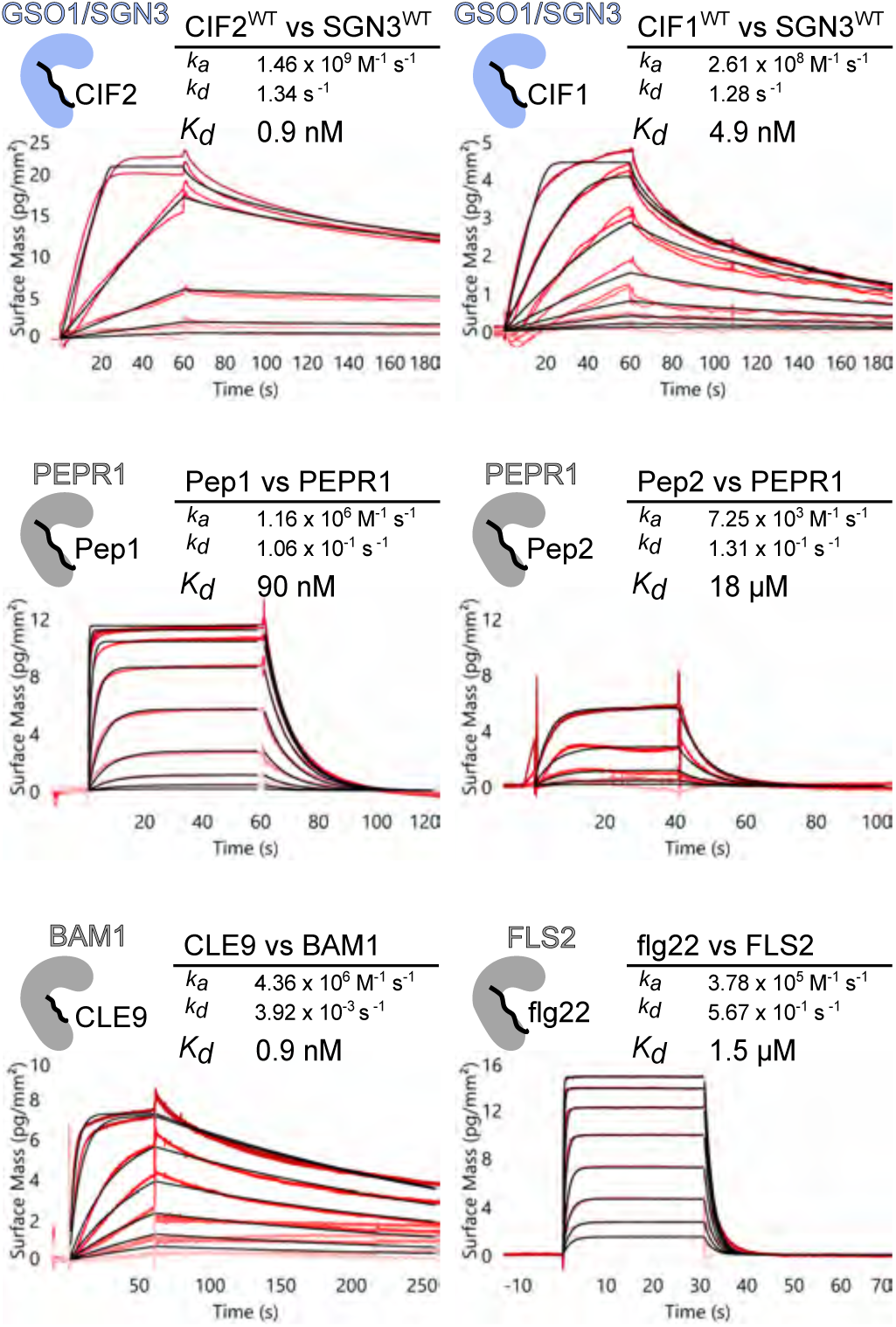
GSO1/SGN3 – CIF2 represents one of the strongest LRR-RK – peptide ligand pairs in Arabidopsis. Quantitative comparison of GSO1/SGN3 – CIF2 with other known peptide ligands binding to their cognate LRR-RKs by grating-coupled interferometry (GCI). Shown are sensorgrams with raw data in red and their respective fits in black. Table summaries of kinetic parameters are shown alongside (ka, association rate constant; kd, dissociation rate constant; Kd, dissociation constant).

To gain mechanistic insight into the GSO1/SGN3 – CIF1/2 interaction, we next determined the crystal structure of a GSO1/SGN3 – CIF2 complex. We produced the GSO1/SGN3 ectodomain (residues 19-870) by secreted expression in insect cells. The native protein did not yield diffraction quality crystals and hence we partially deglycosylated GSO1/SGN3 using a mix of endoglycosidases H, F1 and F3 (see Methods). Crystals obtained in the presence of a synthetic CIF2 peptide diffracted to ~3.0 Å resolution and the structure was solved using the molecular replacement method. The final model contains two GSO1/SGN3 – CIF2 complexes in the asymmetric unit, with a solvent content of ~70 %. The GSO1/SGN3 ectodomain contains 32 LRRs folding into a superhelical assembly previously seen in other plant LRR-RKs (Fig. 2, *SI Appendix*, Fig. S1) (1). The structure completes ~1.5 helical turns, forming the largest LRR ectodomain currently known in plants (Fig. 2). The LRR core is sandwiched between canonical, disulfide bond-stabilized capping domains (Fig. 2, *SI Appendix*, Fig. S1). 16 N-glyosylation sites are evident in the electron density maps of the partially deglycosylated protein, evenly distributed along the spiral-shaped GSO1/SGN3 ectodomain (Fig. 2, *SI Appendix*, Fig. S1). One CIF2 peptide binds in a fully extended conformation to the GSO1/SGN3 LRR core (LRRs 3-23) (Fig. 2, *SI Appendix*, Fig. S1).

**Fig. 2.**
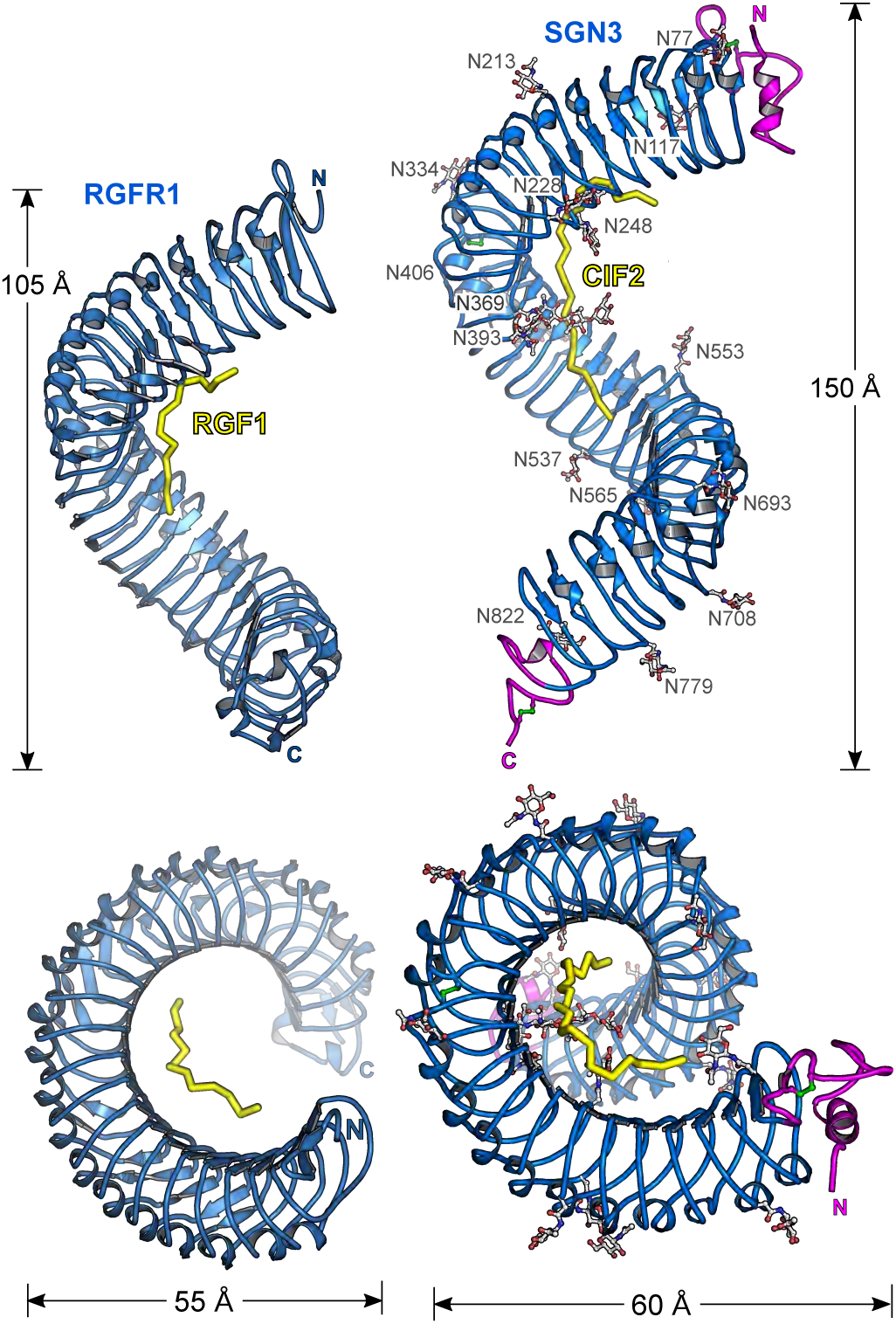
GSO1/SGN3 harbors a large spiral-shaped LRR domain providing the CIF peptide binding surface. Shown is a structural comparison of the SGN3 – CIF2 complex (right) and the RGFR1 – RGF1 complex (left; PDB ID 5hyx, (13)). LRR domains (ribbon diagram) are shown in blue, peptide ligands in yellow (in bonds representation), N- and C- terminal capping domains in magenta, disulfide bonds in green and N-glycans in gray. While the overall architecture and mode of ligand binding is similar in RGFR1 and GSO1/SGN3, the latter receptor contains more LRRs and a much larger peptide binding surface.

We compared our GSO1/SGN3 – CIF2 complex to the previously reported structure of the sTyr-peptide binding receptor RGFR (13). The RGF peptide and the RGFR ectodomain are much smaller compared to CIF2 and GSO1/SGN3 (Fig. 2). However, both RGFR and GSO1/SGN3 provide a binding pocket for the N-terminal sTyr residue and a RxR motif in close proximity to the C-terminus of the respective peptide ligand (*SI Appendix*, Fig. S2). In our structure we find sTyr64 located in a hydrophobic pocket formed by GSO1/SGN3 residues originating from LRRs 3-5 (Fig. 3*A*). It has been previously established that the tyrosylprotein sulfotransferase TPST/SGN2 is genetically required for Casparian strip formation (6). In line with this, recombinant TPST/SGN2 obtained by secreted expression from insect cells has specific tyrosylprotein sulfotransferase activity towards CIF2, using 3’-phosphoadenosine-5’-phosphosulfate as substrate (*SI Appendix*, Fig. S3). The GSO1/SGN3 ectodomain bound tyrosine sulfated CIF2 (CIF2^WT^) with K_*d*_’s of ~2 nM and ~40 nM in GCI and ITC assays, respectively (Fig. 3*B*, *SI Appendix*, Fig. S4). The binding stoichiometry is ~1 in our ITC assays, in agreement with the GSO1/SGN3 – CIF2 complex structure (Fig. 2, *SI Appendix*, Fig. S4). Non-sulfated CIF2^nsY64^ interacted with the GSO1/SGN3 ectodomain with ~100 - 1,000fold reduced binding affinity, depending on the assay used (Fig. 3*B*, *SI Appendix*, Fig. S4). This suggests that the sTyr moiety formed by TPST/SGN2 contributes to the specific recognition of CIF2 by GSO1/SGN3.

**Fig. 3.**
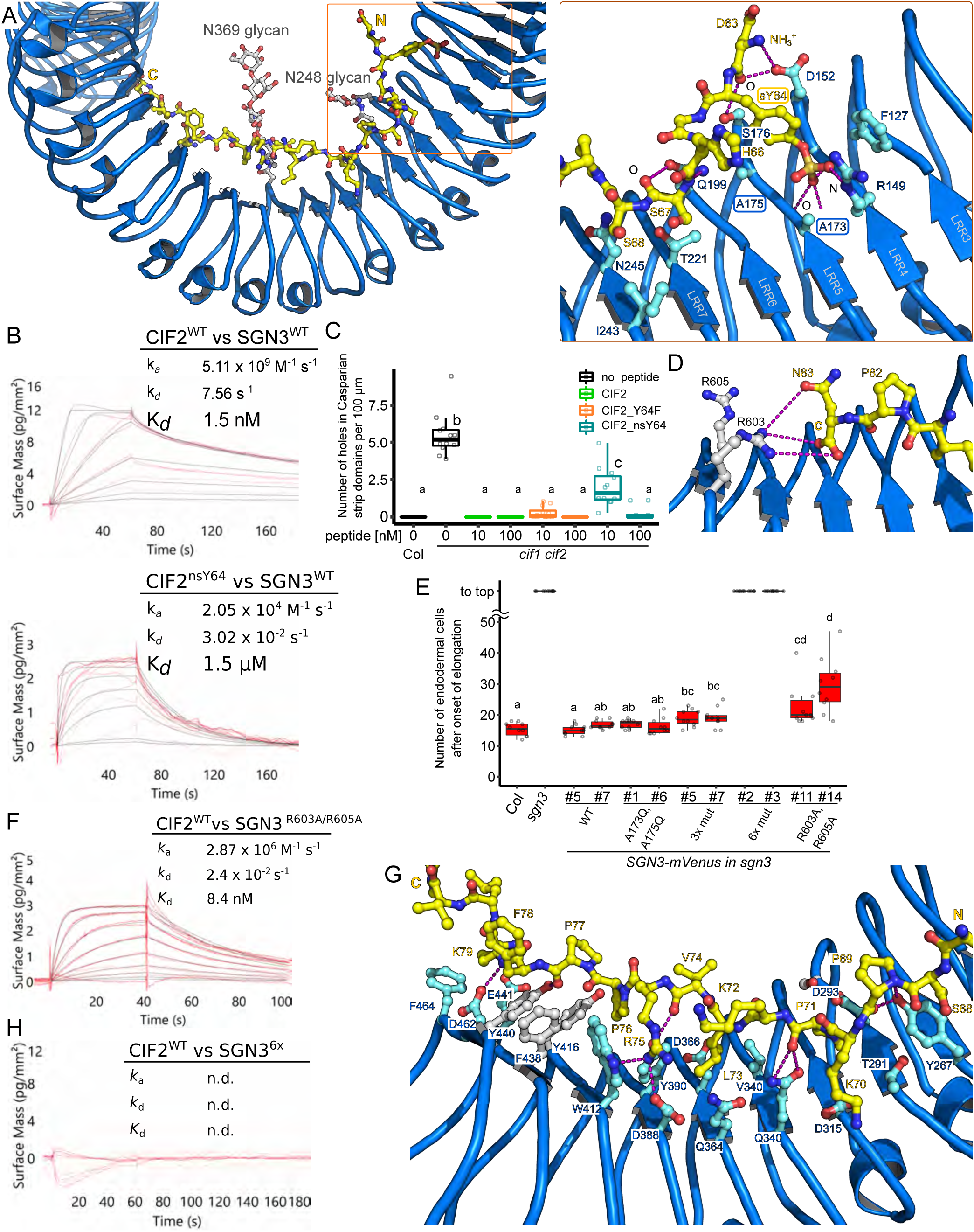
Many peptide – receptor interaction enable high affinity CIF2 binding by GSO1/SGN3. (*A*) (left) Overview of the CIF2 binding site in GSO1/SGN3, colors are as in Fig. 2. (right) Close-up view of the sTyr binding pocket in GSO1/SGN3 with selected residues shown in bonds representation, and with hydrogen bonds indicated as dotted lines (in magenta). (*B*) GCI binding assays of CIF2 variants versus the SGN3 wild-type ectodomain. Raw sensorgrams are shown in red, fitted data in black. Table summaries of kinetic parameters are shown alongside (ka, association rate constant; kd, dissociation rate constant; Kd, dissociation constant). (*C*) Quantitative analyses for the number of holes in Casparian strip domains per 100 μm in *cifl cif2* double mutants with CIF2 peptide-variant treatments (b, c, statistically significant difference with p <0.05, one way ANOVA and Tukey test). (*D*) Close-up view of the GSO1/SGN3 – CIF2 complex. Shown in the C-terminus of the CIF peptide (in bonds representation) and the GSO1/SGN3 RxR motif (in gray). Potential hydrogen bonds are indicated as dotted lines (in magenta) (*E*) Quantification of propidium iodide (PI) staining on *sgn3* mutants complemented with wild-type or mutant SGN3-mVenus under the control of the *SGN3* promoter (no statistically significant difference with one way ANOVA and Tukey test). (*F*) GCI assays of CIF2 versus SGN3 mutant ectodomains. Sensorgrams are shown with raw data in red and their respective fits in black. Table summaries of GCI-derived binding kinetics are shown (ka, association rate constant; kd, dissociation rate constant; Kd, dissociation constant; n.d., no detectable binding). (*G*) Details of the interactions of the CIF2 central part with GSO1/SGN3 LRRs LRRs 6-17. Interface residues are shown in bonds representations, hydrogen bonds as dotted lines (in magenta), amino-acids targeted for the mutational analysis are shown in gray.

To validate our GSO1/SGN3 – CIF2 complex structure, we next replaced the conserved Ala173 and Ala175 from the sTyr binding pocket with glutamine (Fig. 3*A*, *SI Appendix*, Fig. S1). We found that the GSO1/SGN3^A173Q/A175Q^ mutant protein bound CIF2^WT^ and CIF2^nsY64^ with low micromolar affinity in ITC experiments (*SI Appendix*, Fig. S4). In kinetic GCI assays, no specific binding was detected for CIF2^WT^ or CIF2^nsY64^ to GSO1/SGN3^A173Q/A175Q^ (Fig. 3*B*, *SI Appendix*, Fig. S4). However, while removal of the TPST/SGN2-generated sulfation site or mutation of the sTyr binding pocket in the receptor strongly decreased CIF2 binding (~100 – 1,000fold), the non-sulfated CIF2 peptide and the GSO1/SGN3^A173Q/A175Q^ mutant protein complemented *cifl cif2* and *sgn3* loss-of-function phenotypes in Casparian strip formation, respectively (Fig. 3*C,E*, *SI Appendix*, Fig. S5).

We thus analyzed how other amino-acids in the large GSO1/SGN3 CIF2 binding site (~1,500 A^2^ buried surface area) (23) would contribute to the specific recognition of the peptide hormone (Fig. 3*A*). We first mutated the conserved RxR motif in GSO1/SGN3 LRR23, which is involved in the coordination of the C-terminal Asn83 in CIF1/CIF2 (Fig. 3*D*) and in many other plant peptide hormones (1, 7, 13, 16). Replacing Arg603 and/or Arg605 with alanine had a moderate effect on CIF2 binding by GSO1/SGN3 (2-10fold reduction) (Fig. 3F, *SI Appendix*, Fig. S4). In line with this, we find Arg603 and Arg605 not in direct hydrogen bonding distance with either the side-chain of Asn83 or the C-terminal carboxyl group of the CIF2 peptide (Fig. 3*D*). Despite their moderate contribution to CIF2 binding, a GSO1/SGN3^R603A/R605A^ mutant only partially complemented the *sgn3* Casparian strip phenotype (Fig. 3*E*) (see below).

The central part of the CIF peptide binding groove in GSO1/SGN3 is mainly formed by hydrophobic residues and by selected hydrogen bond interactions between residues originating from LRRs 6-17 and backbone atoms from CIF2 (Fig. 3*G*). CIF peptides have been previously demonstrated to be hydroxyprolinated (5) and the corresponding Pro69 and Pro71 residues in CIF2 form part of the central binding site (Fig. 3*G*). While the hydroxyl group of Hyp71 may establish a hydrogen bond with GSO1/SGN3 residue Asp293, we found that CIF2^Hyp69,71^ and CIF2^WT^ bound GSO1/SGN3 with very similar dissociation constants and both could complement the *cifl cif2* Casparian strip phenotype in a same concentration range (*SI Appendix*, Fig. S6).

We replaced three conserved aromatic residues Tyr416, Phe438 and Tyr440 in the central binding groove by alanine (hereafter called SGN3^3x^), and again observed a moderate reduction in CIF2 binding (~10fold) (*SI Appendix*, Fig. S4). Transgenic plants recapitulating these mutations partially rescued the *sgn3* phenotype *in planta* (Fig. 3*E*). However, when we combined this triple mutant with the mutations targeting the sTyr binding pocket in GSO1/SGN3 (SGN3^6x^) (Fig. 3), CIF2 binding was disrupted (Fig. 3*F*, *SI Appendix*, Fig. S4) and the GSO1/SGN3^6x^ mutant failed to complement the *sgn3* phenotype (Fig. 3*E*, *SI Appendix*, Fig. S5). Together, our structural and mutational analysis suggests that GSO1/SGN3 uses a large number of interactions to specifically recognize CIF peptides, requiring numerous receptor – peptide contacts to be altered in order to disrupt CIF peptide binding *in vitro* and GSO1/SGN3 function *in vivo*.

We noted in our structure that outside the sTyr binding pocket, CIF2 mainly uses main-chain atoms to contact the GSO1/SGN3 LRR domain. Thus, sequence-divergent tyrosine sulfated peptides may represent *bona fide* ligands for GSO1/SGN3. Based on this observation, we identified additional, putative CIF peptides in *Arabidopsis* and in other plant species, harboring an N-terminal Asp-Tyr motif required for TPST/SGN2 substrate recognition (10), two central proline residues and a C-terminal His/ Asn residue (*SI Appendix*, Fig. S7). From these candidates we selected the closely related, previously uncharacterized At5G04030 (CIF3 hereafter) and At1G28375 (CIF4) for further analysis (Fig. 4*A*). GCI experiments revealed that tryosine sulfated but not the non-sulfated CIF3 synthetic peptide bound to the GSO1/SGN3 ectodomain with nanomolar affinity (Fig. 4*B*). Due to its hydrophobicity, we could not dissolve the CIF4 peptide in our GCI buffer, and thus performed ITC experiments instead, titrating CIF4 into a GSO1/SGN3 solution containing 5% (v/v) DMSO. In these buffer conditions, CIF4 binds GSO1/SGN3 with 300 nM affinity and with 1:1 binding stoichiometry (Fig. 4*C*). DMSO appears to negatively affect binding, as the CIF2 control bound with ~6fold reduced binding affinity when compared to aqueous buffer conditions (Fig. 4*C*, *SI Appendix*, Fig. S4). Together, the newly identified CIF3 and CIF4 peptides bind to GSO1/SGN3 with high affinity *in vitro*.

**Fig. 4.**
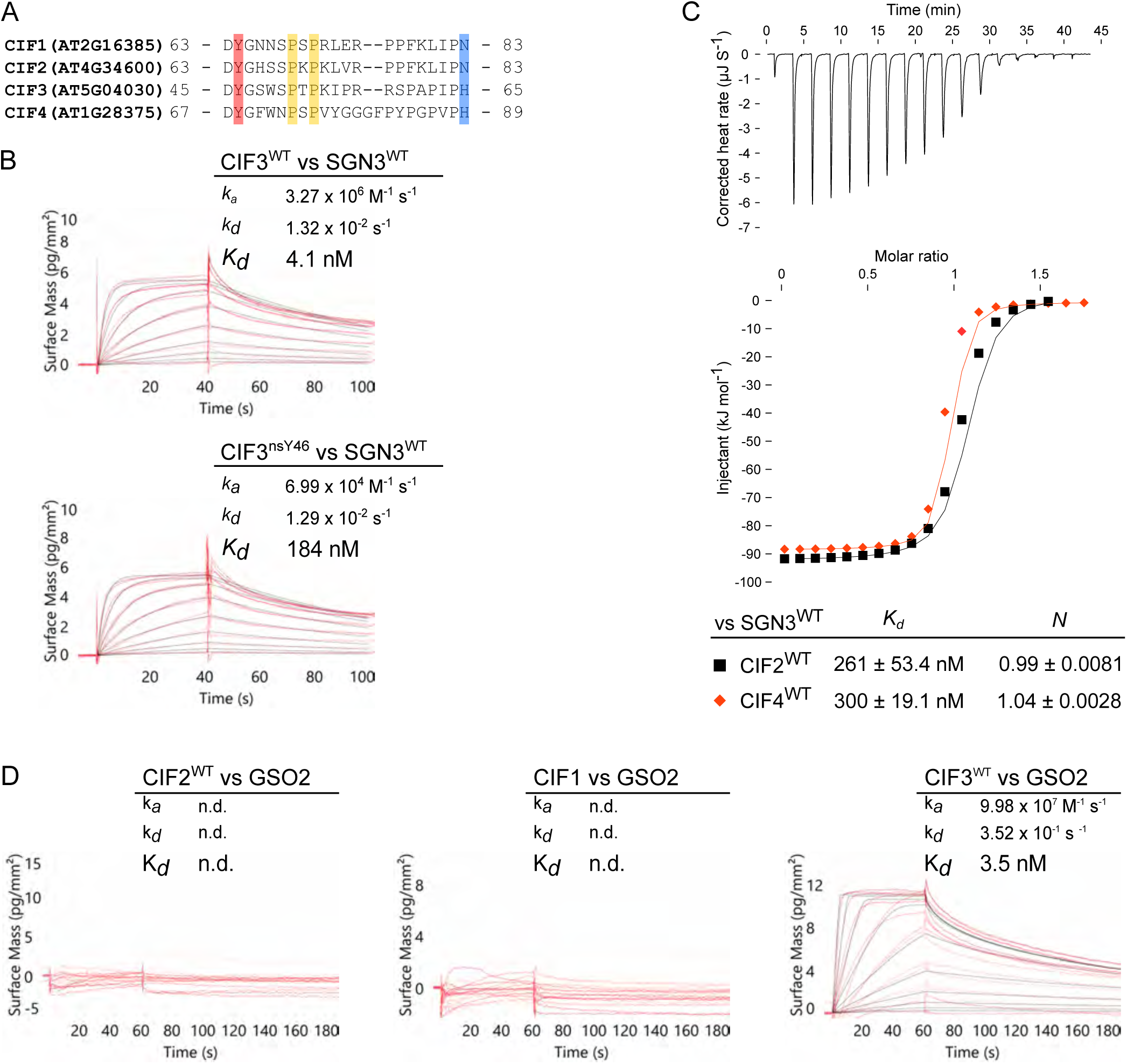
Structure-guided identification of novel CIF peptides. (*A*) Multiple sequence alignment of CIF1 – 4 peptides. The conserved sulfated tyrosine is highlighted in red, hydroxyprolines are in yellow, and the C-terminal asparagine/histidine are shown in blue. (*B*) GCI assays of CIF3 in the presence or absence of sulfation on tyrosine versus the SGN3 wild-type ectodomain. Sensorgrams are presented with raw data in red and their respective fits in black. Table summaries of kinetic parameters are shown alongside (k_a_, association rate constant; k_d_, dissociation rate constant; K_*d*_, dissociation constant). (*C*) ITC assays of CIF2 or CIF4 wild type peptides versus the SGN3 wild type ectodomain. Table summaries for dissociation constants (K*d*) and binding stoichiometries (N) are shown (± fitting error). (*D*) GCI assays of CIF1 – 3 peptides versus the GSO2 wild-type ectodomain.

We next tested if CIFs can also bind to the LRR-RK GSO2, which together with GSO1/SGN3 controls plant embryo development (17). We could purify ~50 μg GSO2 (residues 23-861) from 8 L of insect cell culture, sufficient quantities to perform GCI assays. We found that CIF3 but neither CIF1 or CIF2 bound to the recombinant GSO2 ectodomain (Fig. 4*D*). CIF3 binds both GSO1/SGN3 and GSO2 with a K_*d*_ of ~ 4 nM (Fig. 4*D*). Due to its hydrophobicity, we could not assess binding of CIF4 to GSO2. Together, GSO1/SGN3 and GSO2 display different CIF peptide binding preferences *in vitro*.

In line with our biochemical findings, application of synthetic CIF3 and CIF4 peptides could rescue the *cifl cif2* Casparian strip phenotypes (Fig. 5*A*). However, CIF3 and CIF4 marker lines showed no expression in roots and a *cif3 cif4* double mutant had no apparent Casparian strip or embryo development defect (Fig. 5*B-D*, *SI Appendix*, Fig. S8). Given the fact that we could identify CIF3 and CIF4 orthologs in other plant species (*SI Appendix*, Fig. S7), we speculate these CIF peptides to be involved in yet unidentified GSO1/SGN3 / GSO2 regulated signaling events.

**Fig. 5.**
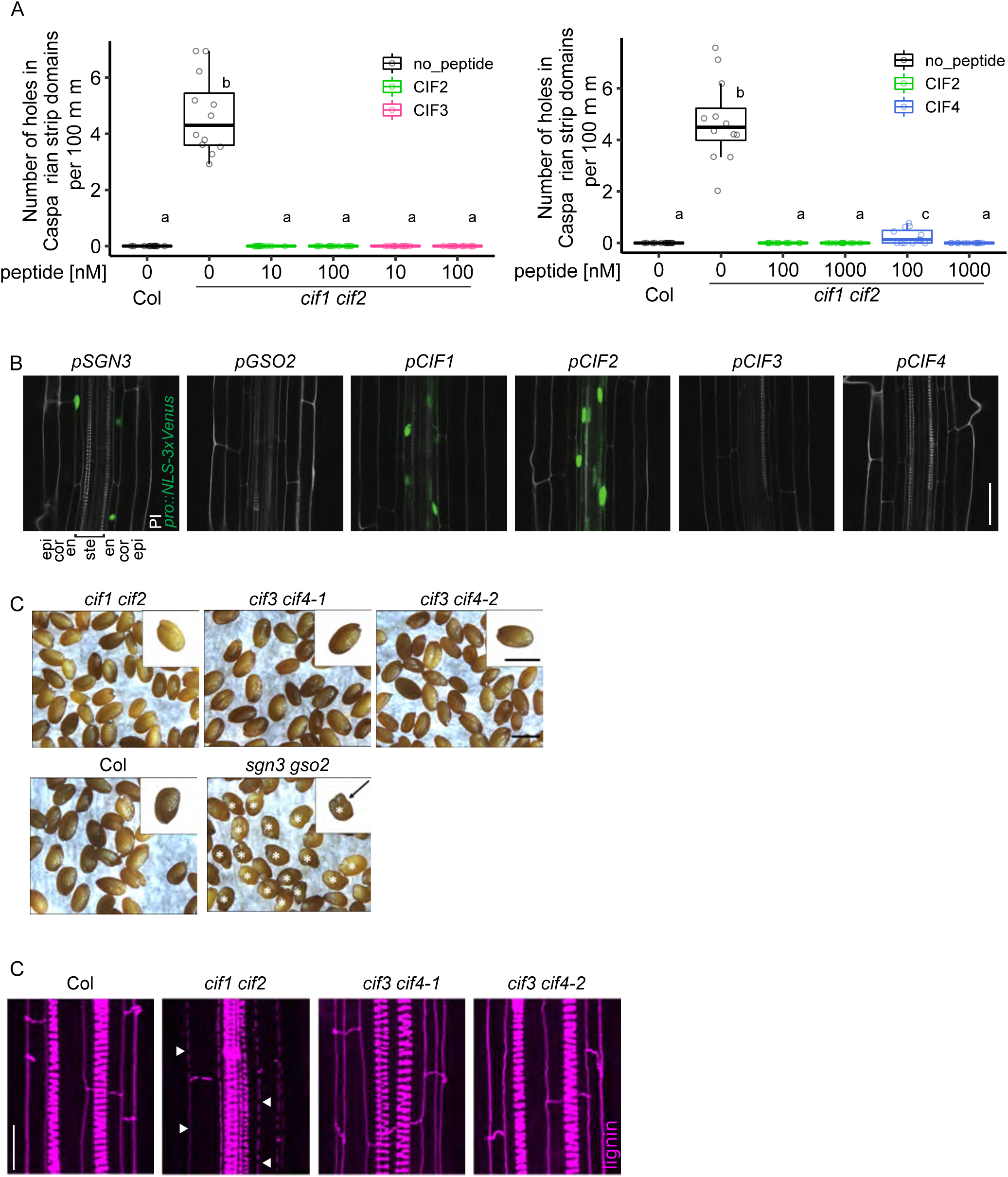
CIF3 and CIF4 are not involved in Casparian strip formation. (*A*) Quantitative analyses of number of holes in Casparian strip domains per 100 μm in Col (WT) or the *cifl cif2* mutant with CIF2, CIF3 or CIF4 peptide treatments (n=12 (experiment with CIF3) and for n≥12 (experiment with CIF4) for each condition). Different letters indicate statistically significant differences (p <0.05, one-way ANOVA and Tukey test). Note that due to the solubility of CIF4, the experiment with CIF4 was done with 0.05% (v/v) DMSO in all conditions including the control. (*B*) Promoter activities around onset of Casparian strip formation. Each promoter drives a NLS (nuclear localization signal)-3xVenus reporter gene. Cell walls were stained with propidium iodide (PI). Cell layers are labeled as Epi (epidermis), Cor (cortex), En (endodermis) and Ste (stele). Scale bar corresponds to 40 μm. (*C*) CIF peptides do not display *gsol gso2* seed shape phenotypes. Show are mature seeds from Col, *ciflcif2*, *cif3 cif4-l*, *cif3 cif4-2* and *sgn3Igsol gso2*. The seeds from *sgn3Igsol gso2* had aberrant shapes (indicated by a *) but seeds from other genotypes showed the normal shapes as did the Col (WT) wild-type control. Scale bars correspond to 0.5 mm. (*D*) *cif3 cif4* double mutants do not show Casparian strip barrier defects. Lignin images were taken around 10 cells after onset of CS. Scale bar corresponds to 20 μm.

Many of the currently known LRR-RKs require the interaction with a shape-complementary co-receptor kinase for high affinity ligand binding and for receptor activation (1, 21). In contrast to, for example, the peptide hormone IDA, CIF1-4 bind to GSO1/SGN3 with nanomolar affinity already in the absence of a co-receptor kinase (Figs. 1,3) (6, 7). This could in principle suggest that GSO1/SGN3 does not require a co-receptor (6). However, we found that both apo and CIF2-bound GSO1/SGN3 ectodomains behaved as monomers in analytical size exclusion chromatography and right-angle light scattering experiments, respectively (Fig. 6*A*). This makes it unlikely that CIF2 binding alters the oligomeric state of GSO1/SGN3, an activation mechanism used by the LRR domain-containing animal Toll-like receptors (24). However, structural features in the GSO1/SGN3 – CIF2 complex suggest that a shape-complementary co-receptor kinase may be required for receptor activation: First, CIF2 contains a C-terminal asparagine residue in close proximity to the GSO1/SGN3 RxR motif (Fig. 3*D*). Both motifs are involved in the recruitment of a SERK co-receptor kinase in the structurally related IDA – HAESA and RGF – RGFR complexes (7, 13). Second, mutation of the RxR motif to alanine has no apparent effect on CIF2 binding *in vitro*, but the mutant receptor can only partially complement the *sgn3* Casparian strip phenotype (Fig. 3*E,F*). Thus, the GSO1/SGN3 RxR motif may not be essential for CIF peptide binding, but may instead be part of a putative receptor – co-receptor complex interface. Third, a surface area covering the C-terminus of the CIF2 peptide and the C-terminal LRRs in GSO1/SGN3 is not masked by carbohydrate, thus representing a potential protein – protein interaction surface (Fig. 6*B*). The corresponding region in SERK-dependent LRR-RKs has been previously shown to represent the receptor – co-receptor complex interface (1).

**Fig. 6.**
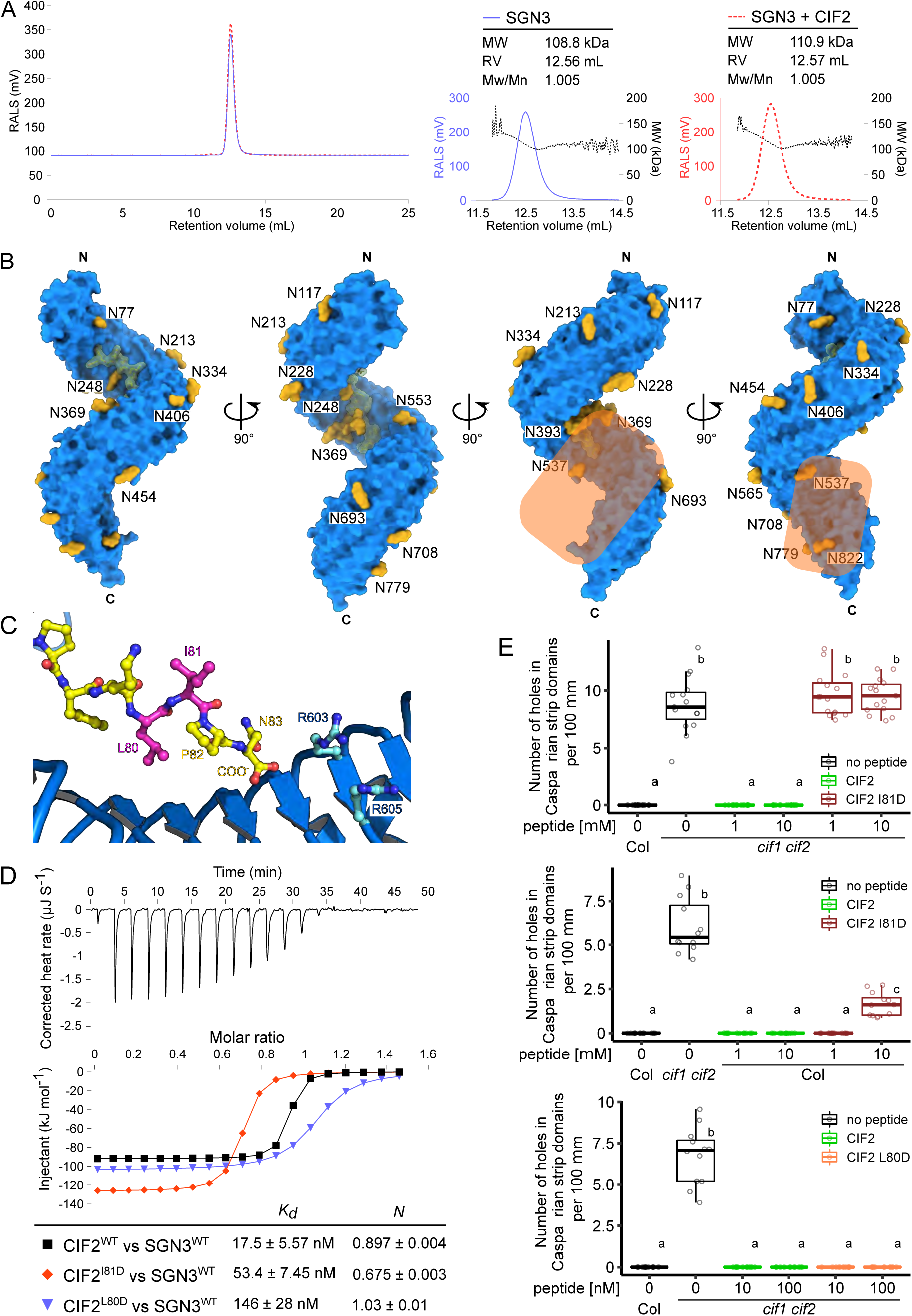
Structural and biochemical evidence for a co-receptor kinase required for GSO1/SGN3 activation. (*A*) Isolated and CIF2-bound GSO1/SGN3 behave as monomers in solution. (*Left*) Analytical size-exclusion chromatography traces of the SGN3 ectodomain in the absence (blue line) or presence (red dotted line) of CIF2 peptides. Right angle light scattering (RALS) traces in the absence (blue) or presence (red) of CIF2 peptides and including the derived molecular masses (black) of GSO1/ SGN3 apo or SGN3-CIF2. Table summaries report the observed molecular weight (MW) and the dispersity (Mw/Mn). The theoretical molecular weight is 94.1 kDa for GSO1/SGN3 (residues 19-870). The GSO1/SGN3 – CIF complex structure reveals a potential co-receptor binding site. Shown is the GSO1/SGN3 ectodomain (surface representation, in blue) in complex with the CIF2 peptide (surface view and bonds representation, in yellow), N-glycans (surface representation in yellow). The potential co-receptor binding surface not masked by carbohydrate is highlighted in orange. (*C*) Close-up view of CIF2 C-terminus bound the GSO1/SGN3, indicating the positions of the side-chains of Leu80 (pointing towards the receptor) and Ile81 (pointing to the solvent) (in magenta). (*D*) ITC assays of CIF2 mutant peptides versus the SGN3 wild type ectodomain. Table summaries for dissociation constants (K_*d*_) and binding stoichiometries (N) are shown (± fitting error). (*E*) Quantitative analyses of number of holes in Casparian strip domains per 100 μm in *cifl cif2* double mutants upon treatment with CIF2 peptide variants. (n=15 for the top panel, n=12 for the middle panel and n≥11 for the bottom panel). Different letters indicate statistically significant differences (p <0.05, one-way ANOVA and Tukey test).

We thus sought to obtain evidence for the involvement of a co-receptor kinase in SGN3 signal transduction. We hypothesized that a co-receptor may bind to the CIF2 C-terminus, coordinated by the GSO1/SGN3 RxR motif (Fig. 6*C*). We replaced CIF2 Ile81, which faces the solvent in our structure, with aspartate (CIF2^I81D^) (Fig. 6*C*) and found that while the mutant peptide still binds GSO1/SGN3 with nanomolar affinity *in vitro* (Fig. 6*D*), it cannot rescue Casparian strip membrane domain formation in *cifl cif2* (Fig. 6*E*). Importantly, wild-type plants treated with micromolar concentrations of CIF2^I81D^ displayed dominant negative Casparian strip integrity phenotypes, while treatment with CIF2^WT^ had no apparent effect (Fig. 6*E*). Mutation of the neighboring Leu80 to aspartate more strongly reduced binding to GSO1/SNG3 when compared to CIF2^I81D^, in agreement with our complex structure, which reveals Leu80 to be part of the CIF2 – GSO1/SGN3 complex interface (Fig. 6*C,D*). CIF2^L80D^ application did not reveal a dominant negative effect but rather rescued the *cifl cif2* double mutant phenotype (Fig. 6*E*). Based on these findings, we speculate that CIF2^I81D^ and CIF2^L80D^ both can bind GSO1/SGN3 *in vivo*, but CIF2^I81D^ specifically blocks interaction with an essential adapter protein required for GSO1/SGN3 activation.

We initially used a reverse genetic approach to identify co-receptors for GSO1/SGN3, based on previous studies on SERKs and SERK-related LRR-RKs (1, 22, 25, 26). However, analysis of known *serk* and *cik*/*nik*/*clerk* loss-of-function mutant combinations revealed no apparent Casparian strip phenotype (*SI Appendix*, Fig. S9). We next performed a biochemical interaction screen, using the known SERK1 and 3 co-receptors as well as other GSO1/SGN3 interacting LRR-RKs, recently identified in a high-throughput biochemical screen (27). From the LRR-RK candidates identified in this screen, we selected putative co-receptors with small LRR ectodomains, including SERK5 (1), CIK/NIK/CLERK proteins recently reported as co-receptors for CLE peptide sensing LRR-RKs (22, 25, 26), the SRF receptor kinases (28), and the immune receptor kinase SOBIR1 (29). We expressed and purified the LRR ectodomains of SERK1, SERK3, SERK5, NIK3, NIK4, SRF3, SRF9 and SOBIR1 and tested for CIF-dependent interaction with the GSO1/SGN3 ectodomain in quantitative GCI assays (Fig. 7*A,B*, *SI Appendix*, Fig. S10). Strikingly, we observed specific binding of SERK1 to GSO1/SGN3 in the presence of either CIF1, 2 or 3, with dissociation constants ranging from ~20 – 300 nM (Fig. 7*C*, *SI Appendix*, Fig. S10). No SERK1 binding to SGN3 was observed in the absence of CIF peptide (*SI Appendix*, Fig. S10), and the co-receptor did not bind the GSO1/SGN3^6x^ mutant (Fig. 7*C*, see above). In line with our structural and physiological assays, the CIF2^I81D^ peptide specifically blocked GSO1/SGN3 – SERK1 interaction, rationalizing its dominant negative effect on Casparian strip formation (Figs. 7*C*). GSO1/SGN3 also interacts with SERK3, but not with SERK5 or any of the other co-receptor candidates derived from the high-throughput screen (*SI Appendix*, Fig. S10) (27). Consistently, we observed specific SERK1/3 binding to GSO2 in the presence of CIF3 (K_*d*_ ~ 20-80 nM) (*SI Appendix*, Fig. S10).

**Fig. 7.**
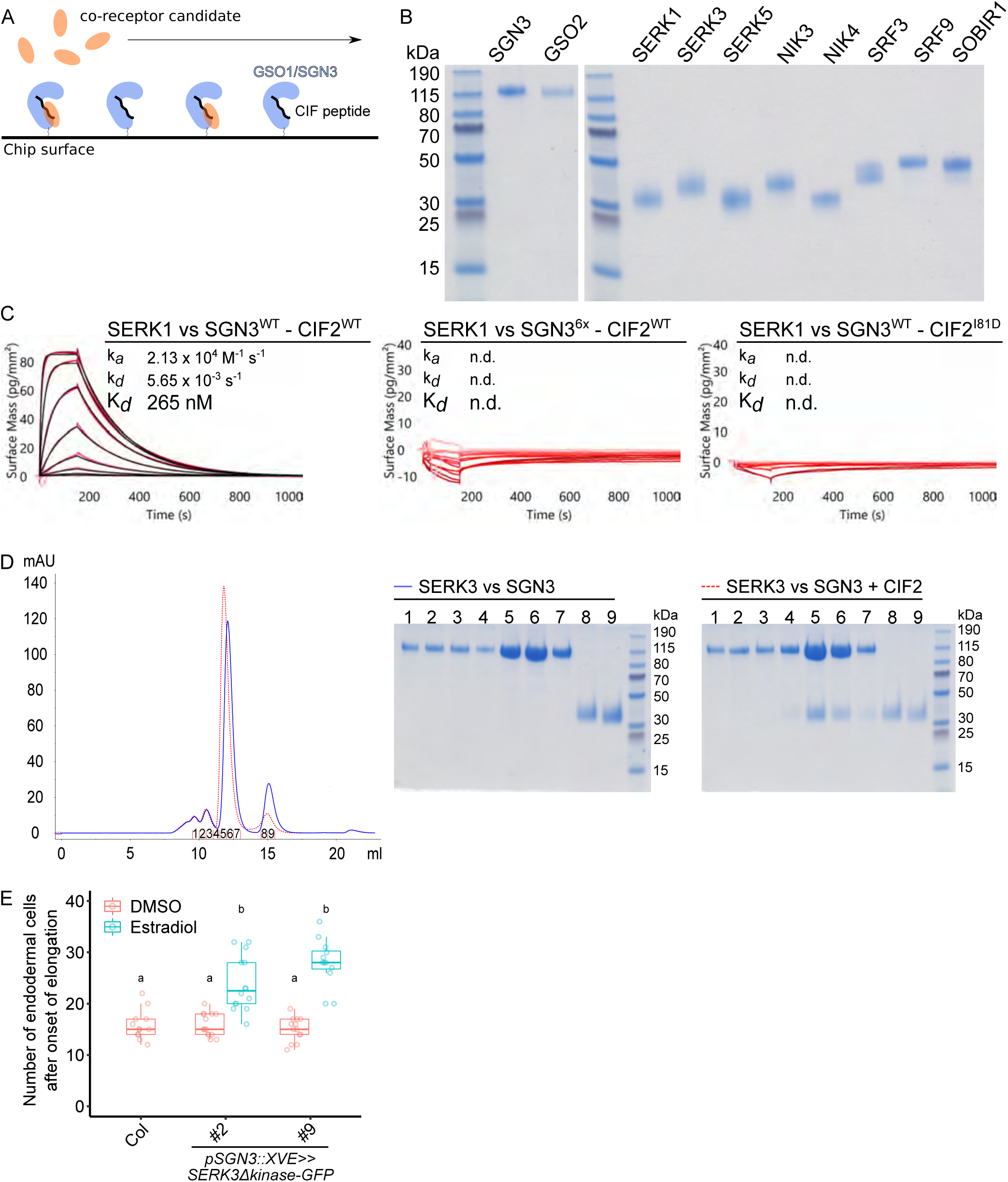
A quantitative interaction screen identifies SERK proteins as putative co-receptors for GSO1/SGN3. (*A*) Schematic overview of the biochemical screen for a GSO1/SGN3 co-receptor. GSO1/SGN3 is immobilized to the GCI chip surface (in blue), the CIF peptide is provided in access in the running buffer (in black) and different recombinantly purified co-receptor candidates are assayed for binding (in orange). (*B*) Coomassie-stained SDS PAGE depicting 1 μg LRR ectodomain of the indicated co-receptor candidate. Shown are isolated monomeric peak fractions from size-exclusion chromatography experiments. (*C*) GCI assays of SERK1 LRR-RK ectodomain versus the SGN3 wild-type and mutant ectodomains in the presence of CIF2 variant peptides. The remaining candidates are shown in *SI Appendix* Fig. S10. Sensorgrams are shown with raw data in red and their respective fits in black. Table summaries of kinetic parameters are shown (k_a_, association rate constant; k_d_, dissociation rate constant; K_*d*_, dissociation constant; n.d., no detectable binding). (*D*) Complex formation of SERK3 and SGN3 ectodomains. (*Left*) Analytical size-exclusion chromatography traces of the SGN3 ectodomain in the absence (blue line) or presence (red dotted line) of CIF2 peptides. An SDS-PAGE analysis of the corresponding fractions is shown alongside. The theoretical molecular weight is 94.1 kDa for SGN3 (residues 19-870) and 21.7 kDa for SERK3 (residues 26 – 220) respectively. (*E*) Induced barrier defect in inducible SERK3 dominant-negative lines. Quantification of barrier permeability was done using the PI assay (n≥12 for each condition). Different letters indicate statistically significant differences (p <0.05, one-way ANOVA and Tukey test).

To our surprise, the interaction of SERKs with ligand-associated GSO1 and GSO2 was much tighter than previously reported for the LRR-RKs BRI1 and HAESA (21). GCI analysis of PEPR1 – Pep1 – SERK1/3 complex formation however revealed an even tighter interaction (K_*d*_’s 1-4 nM), while the related LRR-RK immune receptors FLS2 and EFR bound SERK3 with low micromolar affinity (*SI Appendix*, Fig. S11). Together, our quantitative receptor – co-receptor interaction screen revealed SERK1/3 as *bona fide* co-receptors for GSO1/SGN3 and GSO2. We hypothesized that different SERKs may act redundantly as co-receptor kinases for GSO1/SGN3 in the endodermis, complicating the analysis of *serk* loss-of-function alleles (*SI Appendix*, Fig. S9). We thus generated an estradiol-inducible, dominant-negative SERK3 line (30) and found that it significantly delays Casparian strip formation. While the effect is not as strong as observed for *sgn3* loss-of-function alleles, this provides *in vivo* support for a role of SERK3 and/or SERK homologs in GSO1/SGN3 mediated Casparian strip formation. Taken together, our biochemical and genetic experiments implicate SERK proteins as co-receptors for GSO1/SGN3 and GSO2.

## Discussion

Plants harbor many different classes of signaling peptide hormones, the bioactive forms of which are generated by proteolytic processing from larger pre-proteins and by post-translational modifications including hydroxyprolination and tyrosine sulfation (2). Several of these peptide hormones are specifically sensed by LRR-RKs (1). The 21 amino-acid CIF1 and 2 peptides carry a sulfated tyrosine residue in position 64 *in vivo* (5) and have been shown to represent ligands for the LRR-RK GSO1/SGN3 (5, 6). GSO1/SGN3 tightly interacts with CIF1 and CIF2 with dissociation constants in the low nanomolar range (Fig. 1) (6). The sTyr-containing peptide hormone PSK binds its cognate receptor PSKR with a K_*d*_ of ~1 μM (9). RGF peptides that share the N-terminal Asp-Tyr motif with CIF1/2, interact with different RGFRs with dissociation constants in the high nanomolar to mid-micromolar range (13). Recently, the tyrosine sulfate RaXX peptide from *Xanthomonas* oryzae has been shown to bind the rice LRR-RK XA21 with a Kd of ~ 15 nM (31). Thus, GSO1/SGN3 – CIF1/2 represents the strongest receptor – ligand pair for sTyr-modified signaling peptides currently known in plants. Comparing GSO1/SGN3 – CIF1/2 to known LRR-RK - peptide ligand pairs reveals that plant membrane receptor kinases can sense their cognate peptide ligands with drastically different binding affinities (spanning the micro- to nanomolar range) (Fig. 1) (1, 7). A comparison of the association (k_*a*_) and dissociation rates (k_*d*_) further suggests that high affinity peptide interactions are mainly driven by slow dissociation rates, which however cannot be simply correlated to the size of the respective peptide hormone (Fig. 1). In fact, the 12 amino-acid CLE9 peptide binds the LRR-RK BAM1 with a binding affinity very similar to GSO1/SGN3 – CIF1/2, while the much longer Pep and flg22 peptides bind their cognate receptors with micromolar affinity (Fig. 1). It is of note however that PEPR1 and FLS2 rely on the co-receptor kinase BAK1. BAK1 and other SERK family LRR-RKs have been shown to promote high affinity ligand sensing, with the co-receptor completing the ligand binding pocket and slowing down ligand dissociation (7, 21).

Many plant peptides including the CLE and IDA/IDL families are post-translationally modified, and in both cases these modifications have been shown to be important for high-affinity ligand recognition, and for the bioactivity of the respective peptide hormone (8, 7). For CIF1 and 2, two post-translational modifications have been identified, sulfation of tyrosine 64 and hydroxyprolination of prolines 69 and 71. Using two complementary quantitative binding assays we find that the sulfation of Tyr64 in different CIF peptides is required for high affinity ligand binding to GSO1/SGN3 *in vitro*, but surprisingly removal of the sulfate group from the peptide, or mutation of the sTyr binding pocket in GSO1/SGN3 had little effect on casparian strip formation (Fig. 3). In sharp contrast to for example the HAESA – IDA complex (7), both hydroxyproline residues in CIF2 do not seem to play a major role in ligand sensing, or bioactivity, at least under the conditions tested (*SI Appendix*, Fig. S6). Similarly, the mutation of the GSO1/SGN3 RxR motif conserved among many peptide ligand sensing LRR-RKs (13), had little effect on CIF2 binding and resulted in intermediate Casparian strip formation phenotypes (Fig. 3). We had to go all the way to a GSO1/SGN3 sixtuple mutant to disrupt CIF2 binding *in vitro*, and receptor function *in planta* (Fig. 3). Based on these findings, we speculate that the concentration of mature CIF1 and 2 peptides in the Casparian strip may exceed the nanomolar range, and thus partially functional receptors can still rescue the *sgn3* phenotype. In line with, application of 10-100 nM of non-sulfatable CIF2^Y64F^ can still complement the *cifl cif2* phenotype, despite having a 100 – 1,000fold reduced binding affinity to GSO1/SGN3 (Fig. 3).

Our GSO1/SGN3 – CIF2 structure prompted us to search for additional CIF peptides and we indeed identified several new candidates and characterized CIF3 and CIF4 (Fig. 4, *SI Appendix*, Fig. S7). We found that while GSO1/SGN3 binds CIF1-4 with high affinity, the homologous LRR-RK GSO2 specifically senses CIF3 (Fig. 4). CIF3 and 4 are not expressed in the endodermis (Fig. 5, *SI Appendix*, Fig. S12) and potentially control other, GSO1/SGN3 and GSO2 mediated developmental processes (17, 32). The partially distinct binding specificities of SGN3 and GSO2 suggest that the two receptor have evolved unique functions, possibly to mediate to specific signal inputs in as yet unknown tissue and organ contexts during development. However, a single mutant phenotype for GSO2 has not been described, the only currently known function being redundant with GSO1/SGN3 in embryonic cuticle formation (17). In depth analysis of the GSO2 and CIF3/4 expression domains and targeted phenotyping might identify such a specific, non-redundant function of GSO2 and CIF3/4 in the future. Since neither *cifl cif2*, nor *cif3 cif4* double mutants show an embryonic cuticle phenotype, it will also be important to identify whether a combination of *cifl-4*, possibly a quadruple mutant is required for this developmental process, or whether it is mediated by an additional, thus far unidentified, peptide ligand.

While the high-affinity recognition of CIF peptides by GSO1/SGN3 and GSO2 does not require a co-receptor kinase, the receptor activation mechanism for these LRR-RKs remained to be identified. Despite our initial genetic analyses arguing against a role for the common SERK co-receptor kinases in GSO1/SGN3 function, a quantitative biochemical interaction screen clearly identified SERK1 and 3 as *bona fide* co-receptors. SERKs bind GSO1/SGN3 and GSO2 only in the presence of CIF peptide ligands, suggesting that the previously established ligand-induced receptor – co-receptor heteromerisation mechanism (1, 21) is conserved in GSO1/SGN3 and GSO2 (Fig. 7). CIF3 promotes a much stronger interaction of GSO1/SGN3 or GSO2 with SERK1 when compared to CIF1/2, suggesting that CIF peptides may not only have unique receptor binding specificities, but also different affinities for SERK co-receptors (*SI Appendix*, Fig. S10). It is of note that CIF-dependent interaction of GSO1/SGN3 or GSO2 with SERKs is ~50times stronger than previously described for the LRR-RKs BRI1 and HAESA (21). We speculate that minute amounts of SERK co-receptor may suffice to allow for GSO1/SGN3 receptor activation, possibly rationalizing why *serk* double and triple mutants show no apparent Casparian strip defects (*SI Appendix*, Fig. S9). The dominant negative effect of our SGN3::XVE:SERK3~kinase-GFP line nonetheless provides genetic support for the involvement of SERK proteins in Casparian strip formation (Fig. 7). Generation of clear-cut loss-of-function evidence might prove challenging, since multiple SERK mutants lead to highly pleiotropic phenotypes, including seedling lethality and sterility, in line with their involvement in a large number of LRR kinase-mediated signaling processes (33–35). The biochemical identification of novel CIF peptides and of GSO1/2 co-receptor kinases however now offers new avenues to dissect peptide hormone signaling specificity in a developmental context.

## Acknowledgments

We thank the staff of beam line PXIII of the Swiss Light Source, Villigen, Switzerland for their technical assistance during data collection. This work was supported by grants no. 31003A_176237 and 31CP30_180213 (M.H.), 31003A_156261 and 310030E_176090 (N.G.) from the Swiss National Science Foundation, an ERC Consolidator Grant (616228-ENDOFUN) (N.G.), a Human Frontier Science Program Organization (HFSPO) postdoctoral fellowship no. LT000567/2016-L (S.O.), a Japanese Society for the Advancement of Science (JSPS) fellowship (S.F.), and by an International Research Scholar Award from the Howard Hughes Medical Institute (M.H.).

## Materials and Methods

### Protein expression and purification

SGN3 (residues 19 - 870) coding sequence was amplified from the AP018 plasmid containing SGN3 cDNA (19). GSO2 (residues 23 - 861), TPST (residues 25 - 441), SERK1 (residues 24 - 213), SERK3 (residues 1 – 220), NIK3 (residues 26 – 238), NIK4 (residues 31 – 238), SRF3 (residues 1 – 316), and SRF9 (residues 1 – 334) were amplified from *A. thaliana* cDNA, SOBIR1 (residues 1 - 270), PEPR (residues 1 - 767), FLS2 (residues 1 – 800), and EFR (residues 1 - 642) from *A. thaliana* genomic DNA. BAM1 (residues 20 – 637), and SERK5 (residues 24 - 214) were synthesized (Geneart, Germany) with codons optimized for expression in *Trichoplusia ni*. The constructs were cloned in a modified pFastBac vector (Geneva Biotech) containing an azurocidin signal peptide, except for SERK2, SERK3, SRF3, SRF9, SOBIR1, PEPR, and FLS2 with a native secretion signal peptide, respectively, and a TEV (tabacco etch virus protease) cleavable C-terminal StrepII – 9x His tag. SGN3 and GSO2 were also cloned into the vector harboring the *Drosophila* BiP secretion signal peptide, which was amplified from B02_SRF6_pECIA2 (27), a C-terminal TEV cleavable StrepII – 10x His tag and a non-cleavable Avi-tag (36, 37). SGN3 variants carrying point mutations were generated using the primer extension method for site-directed mutagenesis. *Trichoplusia ni* (strain Tnao38) (38) cells were infected with a multiplicity of infection (MOI) of 1 at a density of 2 × 10^6^ cells ml^−1^ and incubated for 26 h at 28 °C and for additional 48 h at 22 °C. The secreted protein was purified from the supernatant by Ni^2+^ (HisTrap Excel; GE healthcare; equilibrated in 50 mM KP_i_ pH 7.6, 250 mM NaCl, 1 mM 2-Mercaptoethanol) and StrepII (Strep-Tactin XT Superflow high affinity chromatography: IBA; equilibrated in 20 mM Tris pH 8.0, 250 mM NaCl, 1 mM EDTA) affinity chromatography. The tag was cleaved with His-tagged TEV protease at 4 °C overnight and removed by a second Ni^2+^ affinity chromatography step. Proteins were then further purified by size-exclusion chromatography on either a Superdex 200 increase 10/300 GL, Hi Load 16/600 Superdex 200 pg, or HiLoad 26/600 pg column (GE Healthcare), equilibrated in 20 mM sodium citrate pH 5.0, 250 mM NaCl. For crystallization, the SGN3 protein was dialyzed against 20 mM sodium citrate pH 5.0, 150 mM NaCl and treated with Endoglycosidase H, F1, and F3 to trim N-glycan chains, followed by size-exclusion chromatography to further purify the deglycosylated SGN3. His-tagged BirA was purified from *E. coli* by Ni^2+^ affinity chromatography.

### Crystallization and data collection

Crystals of the deglycosylated SGN3 in complex with the CIF2 peptide developed at room temperature in hanging drops composed of 1 μl protein solution (1 mg ml^−1^) containing 0.5 mM CIF2 and 1 μl of crystallization buffer (17 % [w/v] PEG 6,000, 0.1 M Tris pH 7.5, 0.2 M LiCl), suspended above 1.0 ml of the latter as reservoir solution and using microseeding protocols. Crystals of SGN3 in complex with the CIF2^Hyp69, 71^ peptide developed in crystallization buffer (16 % [w/v] PEG 4,000, 0.1 M Tris pH 8.5, 0.2 M MgCl_2_). Crystals were cryo-protected by serial transfer into crystallization buffer supplemented with 20 % (v/v) glycerol (SGN3 – CIF2) or 20 % (v/v) ethylene glycol (SGN3 – CIF2^Hyp69, 71^) and cryo-cooled in liquid nitrogen. Sulfur single-wavelength anomalous diffraction (SAD) data to 4.0 Å resolution was collected at beam-line PXIII at the Swiss Light Source (SLS), Villigen, CH with λ = 2.066 Å. A native data set to 2.95 Å resolution was collected on a crystal from the same drop cryo-protected by same way with λ = 1.0 Å. Data processing and scaling was done in XDS (39).

### Structure solution and refinement

The structure was solved using the molecular replacement method as implemented in the program PHASER (40), and using the isolated ectodomain of the LRR-RK PEPR as search model (PDB-ID 5gr8). The solution comprised a dimer in the asymmetric unit and the structure was completed in alternative cycles of manual model building in COOT (41) and restrained TLS refinement in phenix.refine (42). A phased anomalous difference electron density map calculated with the program ANODE (43) was used to assign the position of disulfide bonds and free cysteines/methionines in the the structure. Analysis with phenix.molprobity (44) reveal good stereochemistry of the final model. Structural diagrams were prepared using Pymol (https://sourceforge.net/projects/pymol/) and povray (http://www.povray.org/).

### Grating – coupled interferometry

GCI experiments were performed with the Creoptix WAVE system (Creoptix AG, Switzerland) using either 4PCP or 4PCH WAVE chips (thin quasiplanar polycarboxylate surface or quasiplanar polycarboxylate surface with high capacity, respectively; Creoptix, Switzerland). For direct amine coupling, chips were conditioned with borate buffer (100 mM sodium borate pH 9.0, 1 M NaCl; Xantec, Germany) and the respective ligands were immobilized on the chip surface using standard amine-coupling; 7 min activation (1:1 mix of 400 mM *N*-(3-dimethylaminopropyl)-*N’*-ethylcarbodiimide hydrochloride and 100 mM *N*-hydroxysuccinimide (Xantec, Germany)), followed by injection of the ligands (50 – 100 μg ml^−1^) in 10 mM sodium acetate pH 5.0 (Sigma, Germany) until the desired density was reached, passivation of the surface (0.5 % BSA (Roche, Switzerland) in 10 mM sodium acetate pH 5.0) and final quenching with 1M ethanolamine pH 8.0 for 7 min (Xantec, Germany). For biotinylated ligands capturing, streptavidin (50 μg ml^−1^; Sigma, Germany) was immobilized on the chip surfaces with same method with the direct amine coupling, followed by capturing respective biotinylated ligands (50 – 100 μg ml^−1^) until the desired density was reached. Kinetic analyses for peptide ligands were performed at 25°C with a 1:2 dilution series from 100 nM for CIF variants in the presence of sulfation or 10 μM in the absence of sulfation, for a co-receptor screen using the biotinylated ligands-captured chips with a 1:3 dilution series from 6.7 μM for SERK1, 3 or 20 μM for the others in 20 mM citrate pH 5.0, 250 mM NaCl, 0.01 % Tween 20. Blank injections were used for double referencing and a dimethylsulfoxide (DMSO) calibration curve for bulk correction. Analysis and correction of the obtained data was performed using the Creoptix WAVE control software (correction applied: X and Y offset; DMSO calibration; double referencing). Mass transport binding models with bulk correction were used for the experiments of SGN3 – CIF peptides binding and one-to-one binding models for the other experiments.

### Isothermal titration calorimetry

All ITC experiments were perfomed on a MicroCal PEAQ-ITC (Malvern Panalytical) with a 200 μl sample cell and a 40 μl injection syringe at 25 °C. Proteins were dialyzed into ITC buffer (20 mM sodium citrate pH 5.0, 250 mM NaCl, exceptionally containing 5 % (v/v) DMSO for CIF4 experiments) prior to all titrations. A typical experiment consisted of injecting 200 μM CIF peptide in 2 μl intervals into the cell containing 20 μM GSO1/SGN3 receptor. The MicroCal PEAQ-ITC analysis software (version 1.21) was used for data analysis.

### Right-angle light scattering

The oligmeric state of SGN3 was analyzed by size exclusion chromatography with a right angle light scattering (RALS), using an OMNISEC RESOLVE / REVEAL combined system (Malvern Panalytical). Instrument calibration was performed with a BSA standard (Thermo Scientific Albumin Standard). 20 μM SGN3 in the presence or absence of 100 μM CIF2, in a volume of 50 μl, were separated on a Superdex 200 increase 10/300 GL column (GE Healthcare) in 20 mM sodium citrate pH 5.0, 250 mM NaCl, at a column temperature of 35 °C and a flow rate of 0.7 ml min^−1^. Data were analyzed using the OMNISEC software (version 10.41).

### Biotinylation of proteins

The respective proteins (20 – 100 μM) were biotinylated with biotin ligase BirA (2 μM) (37) for 1 h at 25 °C, in a volume of 200 μl; 25 mM Tris pH 8, 150 mM NaCl, 5 mM MgCl2, 2 mM 2-Mercaptoethanol, 0.15 mM Biotin, 2 mM ATP, followed by size-exclusion chromatography to purify the biotinylated proteins.

### Sulfotransferase assay

Sulfotransferase assays were performed with universal sulfotransferase activity kit (R&D systems, UK). Non-sulfated CIF2 (residues 59 – 72) (1 mM) were mixed with TPST using a 1:2 dilution series from 1 μM (48 ng μl^−1^) in a volume of 50 μl; 50 mM Tris pH 7.5, 50 mM NaCl, 15 mM MgCl_2_, 0.2 mM 3’-Phosphoadenosine 5’-phosphosulfate (PAPS), phosphatase (500 ng) for 30 min at 30 °C. 30 μl of malachite green reagent A and B, 100 μl of distilled water was added to each sample and incubated for 20 min at 30 °C. The absorption of each sample at 620 nm was determined with a microplate reader (Synergy2, Biotek). Phosphate standard curves were determined using a 1:2 dilution series starting from 100 mM KH_2_PO_4_. Product formation was calculated using the conversion factor from the phosphate standard curve.

### Analytical size-exclusion chromatography

Gel filtration experiments were performed using a Superdex 200 Increase 10/300 GL column (GE Healthcare) equilibrated in 20 mM sodium citrate pH 5.0, 250 mM NaCl. A 500 μl aliquot of SGN3 and SERK3 (at a concentration of 10 μM) was loaded sequentially onto the column and elution at 0.75 ml min^−1^ was monitored by ultraviolet absorbance at 280 nm. The CIF2 peptide concentration was 20 μM in the SGN3 – CIF2 – SERK3 complex sample prior to loading.

### Plant material and growth conditions

For all experiments, *Arabidopsis thaliana* (ecotype Columbia) was used. T-DNA tagged lines for *sgn3-3* (SALK_043282), *gso2* (SALK_143123C) and *cif3-2* (GABI_516E10) were obtained from NASC (http://arabidopsis.info/) and GABI (https://www.gabi-kat.de/) respectively. The *cifl-2 cif2-2* double mutant and *cif4* mutant were generated by CRISPR-Cas9 technique in Col wildtype or *cif3-2* mutant background (see below). Insertion points of the T-DNA and the CRISPR lines were verified by Sanger sequencing. Plants were grown on half-strength Murashige-Skoog (MS) agar (1%) for 5d vertically after 2d stratification at 4°C in the dark. For peptide (Peptide Specialty Laboratories GmbH) treatment assays, seeds were germinated on medium with or without the indicated peptide concentrations and grown for 5d. Estradiol (Sigma) was dissolved in DMSO and used at 5 μM final concentration. DMSO concentration was 0.05% (v/v) at final dilution.

### Molecular cloning

For promoter reporter lines, upstream regions of each gene - indicated by ‘length upstream of ATGs’ - were cloned into gateway entry vectors and fused to NLS-3 x Venus via an LR reaction (pSGN3 5583 bp, pGSO2 3893 bp, pCIF1 1797 bp, pCIF2 1756 bp, pCIF3 2092 bp and pCIF4 2201bp). The pSGN3::SGN3-mVenus construct (19) was used as template to generate SGN3-mVenus variants by site-directed mutagenesis. CRISPR-Cas9 constructs were generated following a published method (45) after switching selection markers from Basta to FASTRed in the final construct with *S. pyogenes* Cas9. For generating *cifl-2* and *cif2-2*, 5’-ttgggtataagcttgaaagg -3’ and for generating *cif4-l* and *cif4-2*, 5’-aacccaagcccggtttacgg -3’ and 5’-ttggatttcaccctaaacga -3’ primers were used respectively. For constructing the dominant negative SERK3 (pSGN3::XVE≫SERK3(residues 1-243)-GFP), a fragment of SERK3 genomic region (residues 1-243.) was cloned into an entry vector and fused with pSGN3::XVE-LexA and GFP via a LR reaction. The constructs were transformed into the wild-type or *sgn3* mutant plants using the *Agrobacterium tumefaciens* GV3101 (MP90)-mediated floral dip method (46).

### Microscopy

Signals were visualized using an SP8 microscope (Leica). Excitation and detection windows, respectively, were as follows: GFP (488 nm, 500-550 nm), Venus or mVenus (514 nm, 520 – 580 nm), propidium iodide (488 nm, 600 – 650 nm) and fuchsin (561 nm, 570 – 650 nm). Images were processed using the Fiji package of ImageJ (47).

### Propidium iodide barrier assay

5d old seedlings were incubated in 10 μg/mL propidium iodide (PI) - water solution for 10 min and transferred into fresh water. For quantification, “onset of cell elongation” was defined as the point where endodermal cell length exceeded two times its width in a median longitudinal section. Cell counting was done using a Zeiss LSM 700 with a 488 nm laser and an SP640 filter split at 600 nm.

### Visualization of lignin

Lignin staining was performed as described in previous reports (48, 49). Briefly, 5d old seedlings were fixed in 4% (v/v) paraformaldehyde PBS solution (pH 6.9) for 1h without vacuum treatment. The samples were rinsed with PBS twice and incubated in ClearSee (10% (w/v) xylitol, 15% (w/v) sodium deoxycholate, 25% (w/v) urea in water) solution overnight. After removing the solution, samples were stained with 0.2% fuchsin in ClearSee solution overnight. Fuchsin solution was removed and the seedlings were briefly rinsed with fresh ClearSee solution and washed by gently agitation in fresh ClearSee solution for 30 min. After exchanging the ClearSee solution, the seedlings were washed overnight.

---

**Fig. S1.**
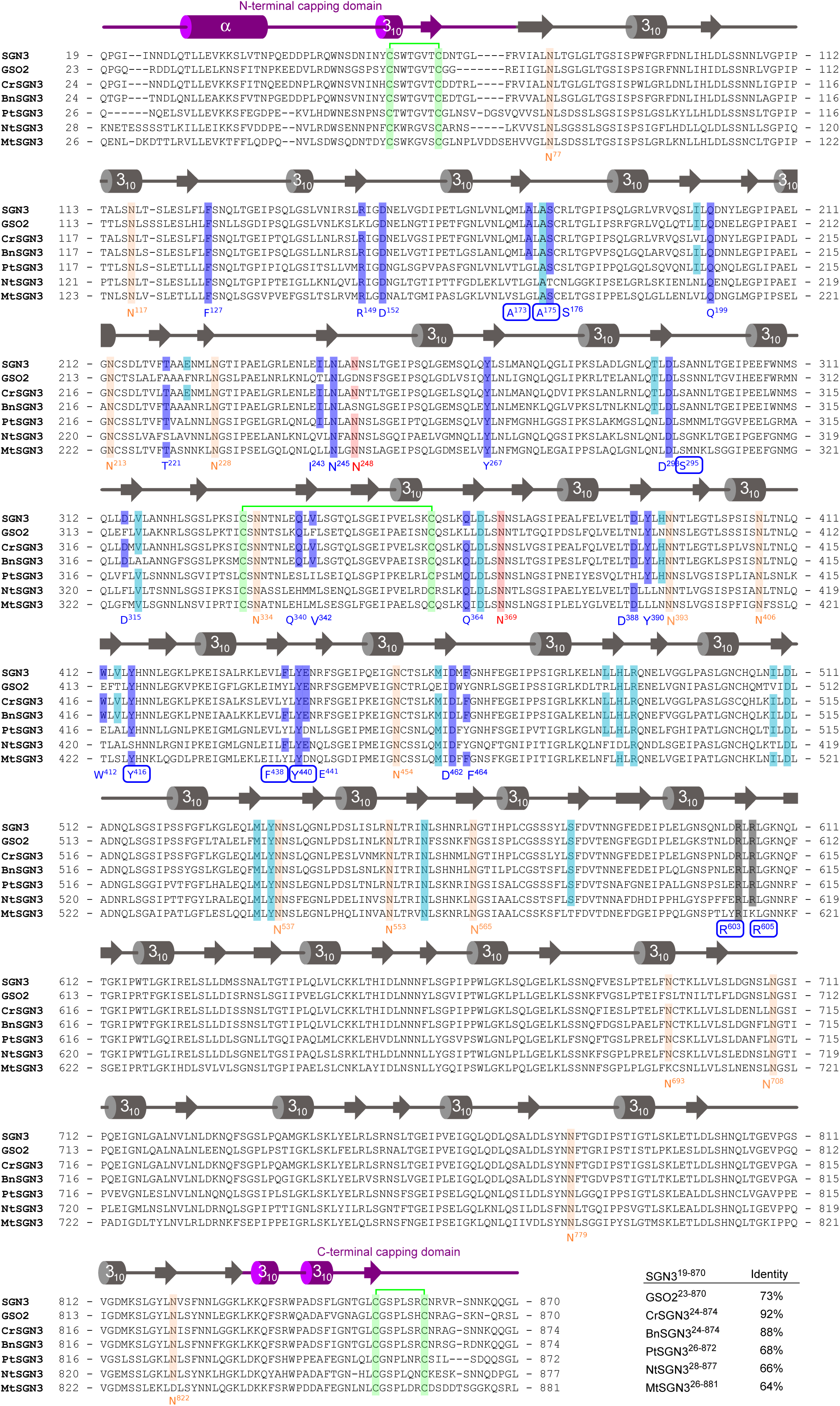
Structure-based multiple sequences alignment of SGN3 ectodomains from *Arabidopsis thaniana*. GSO1/SGN3 (NCBI (https://www.ncbi.nlm.nih.gov/) identifier: OAO97463), GSO2 (NCBI identifier: OAO90459), *Capsella rubella* SGN3 (NCBI identifier: XP_006285037.2), *Brassica napus* SGN3 (NCBI identifier: XP_013660918.1), *Populus trichocarpa* SGN3 (NCBI identifier: XP_002299384.1), *Nicotiana tabacum* SGN3 (NCBI identifier: XP_016509707.1), and *Medicago truncatula* SGN3 (NCBI identifier: XP_013457406.1). A secondary structure assignment, calculated with DSSP (50), is shown beside. SGN3 residues forming hydrogen bonds with CIF2 in the SGN3 – CIF2 complex are highlighted in blue, residues interacting with CIF2 in cyan, glycosylated asparagine residues in orange, asparagine residues with glycans directly contacted with CIF2 in red, RxR motif in gray, cysteines forming disulfide bonds in light green. All numbering refers to AtSGN3. Table summarizes amino acid sequence identities among SGN3 ectodomains versus AtSGN3.

**Fig. S2.**
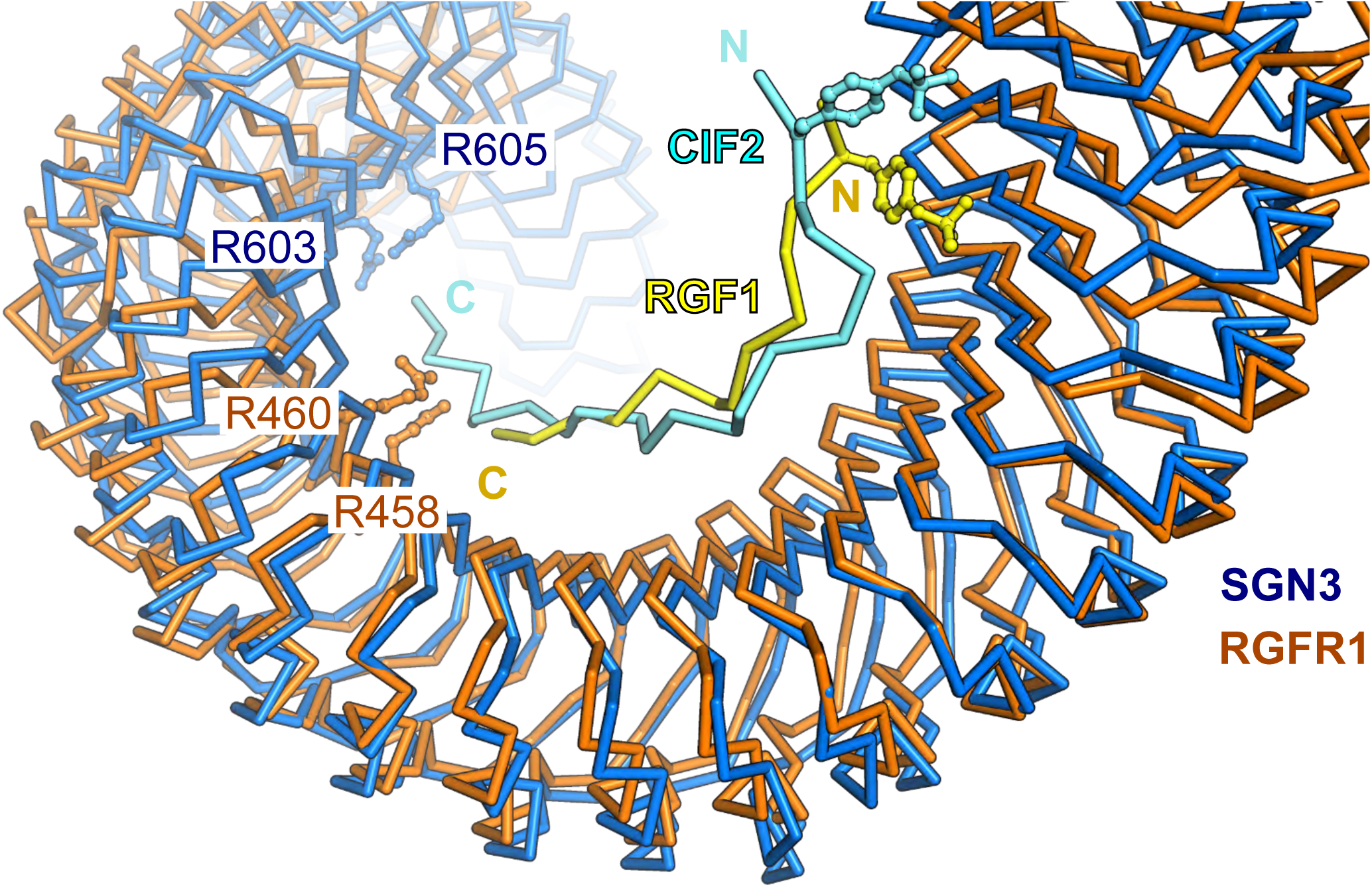
Different LRR-RKs binding tyrosine sulfated peptide share structural similarity. Structural superposition of SGN3 – CIF2 (blue and cyan, respectively) and RGFR – RGF1 (orange and yellow; PDB ID 5hyx) complex structures. Asparagine residues of the RxR motif are shown. The two complex structures align with a root mean square displacement (r.m.s.d.) ~ 3.1 Å comparing 498 corresponding C_a_ atoms.

**Fig. S3.**
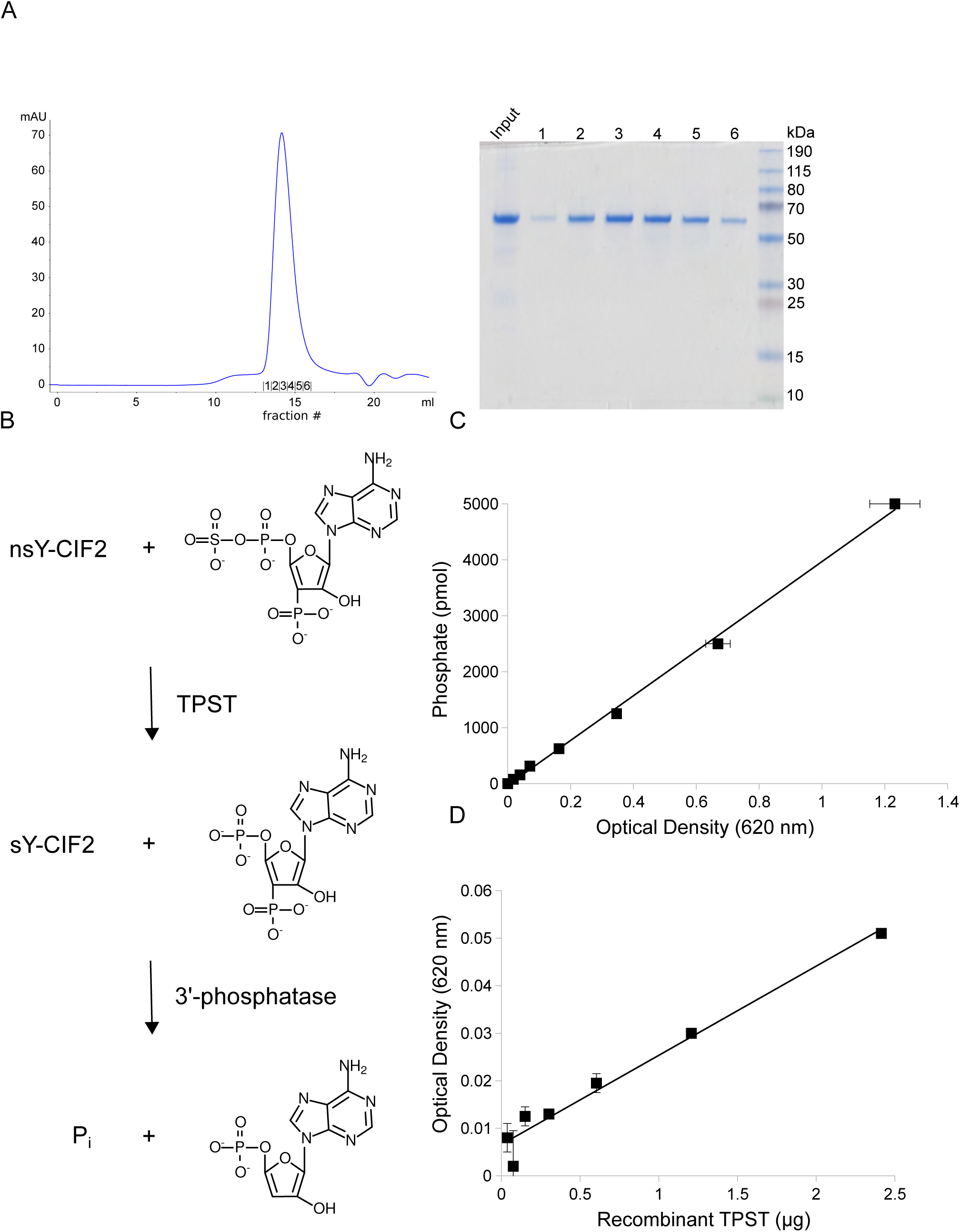
CIF2 is a substrate of the plant tyrosylprotein sulfotransferase TPST/SGN2. (*A*) Size-exclusion chromatography trace of TPST (residues 25 – 441) purified from insect cells. (*Right*) Coomassie-stained SDS PAGE of the corresponding elution fractions. (*B*) Scheme of sulfotransferase assays. Inorganic phosphate (Pi) release was detected using a malachite green Pi quantification assay to calculate the kinetics of the sulfotransferase reaction. (*C*) Pi standard curve used for the enzymatic assay. (*D*) 0.2 mM 3’-Phosphoadenosine 5’-phosphosulfate (PAPS) was incubated with varying concentrations of TPST enzyme for 30 min at 30 °C. Optical densities (ODs) were plotted versus the amount of TPST recombinant protein. A specific activity (1.25 pmol min^−1^ μg^−1^) was calculated.

**Fig. S4.**
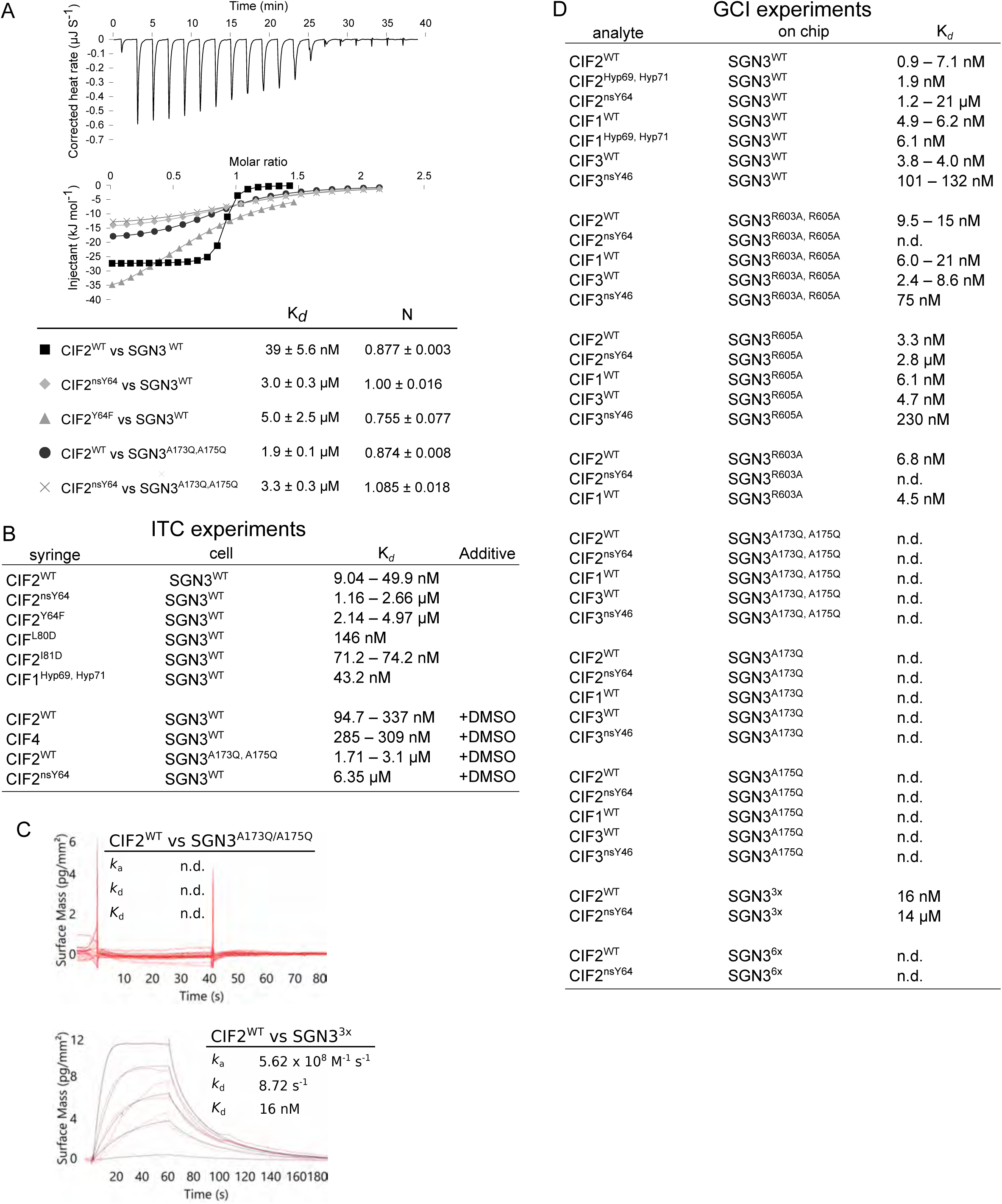
Mutational characterization of the GSO1/SGN3 – CIF2 complex interface. (*A*,*B*) Isothermal titration calorimetry (ITC) assays of CIF2 variants versus SGN3 wild-type and mutant ectodomains. Table summaries for dissociation constants (K*d*) and binding stoichiometries (N) are shown (± fitting error). (*C*,*D*) GCI assays of CIF variants versus SGN3 wild-type and mutant ectodomains. sensorgrams are represented with raw data in red and their respective fits in black. Table summaries of kinetic parameters are shown alongside (k_a_, association rate constant; k_d_, dissociation rate constant; K_*d*_, dissociation constant; n.d., no detectable binding).

**Fig. S5.**
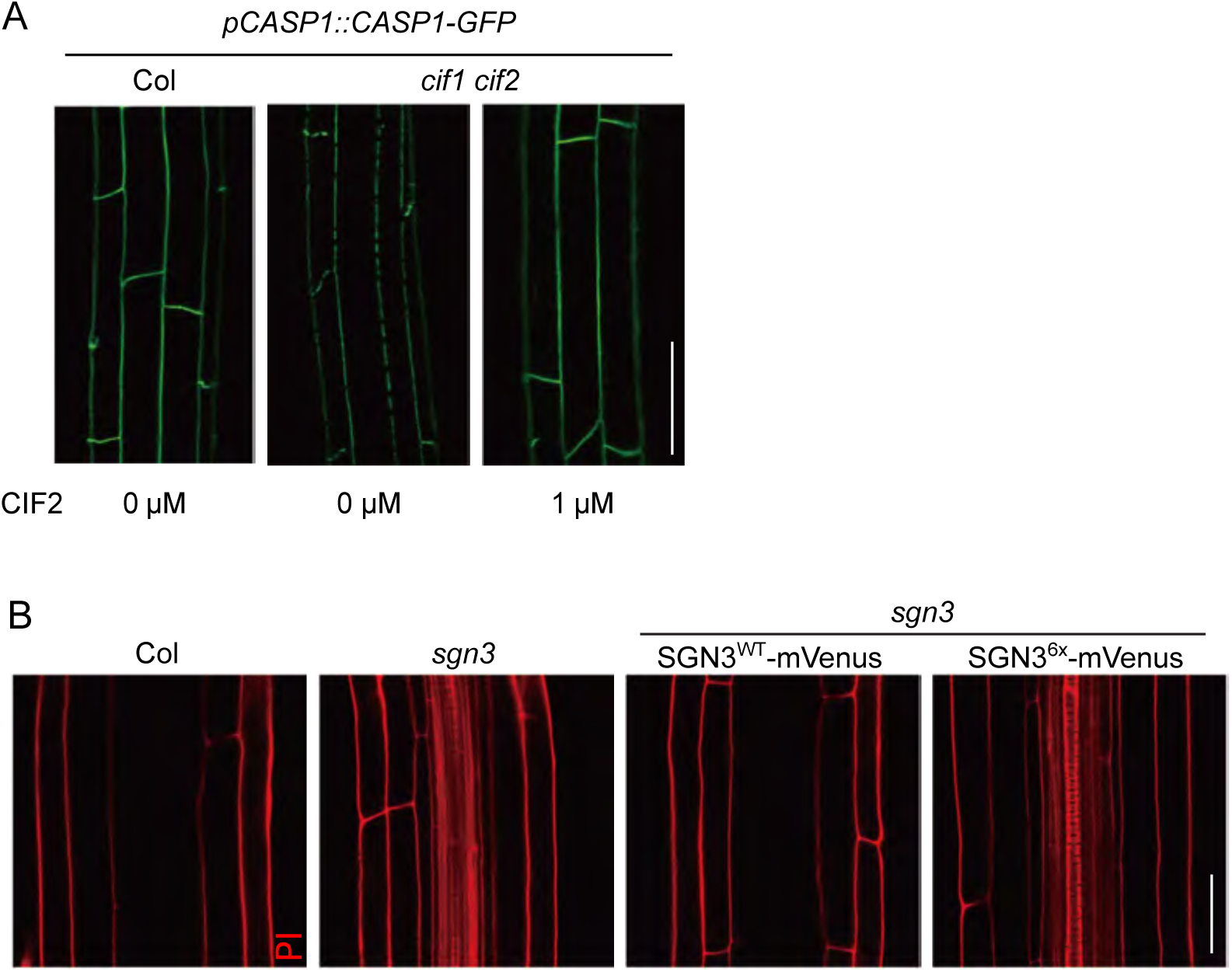
The GSO1/SGN3 6x mutant fails to complement the *sgn3* Casparian strip phenotype. Casparian strip domains are visualized in Col (WT) and *cifl cif2* with or without CIF2. Scale bar = 20 μm (*B*) Representative images of PI permeability in the roots of the indicated genotypes. Pictures were taken around 25-30 cells after onset of endodermal cell elongation. *sgn3* and *sgn3* transformed with SGN36x-mVenus both display staining of vasculature, indicative of barrier defect. Scale bar = 40 μm.

**Fig. S6.**
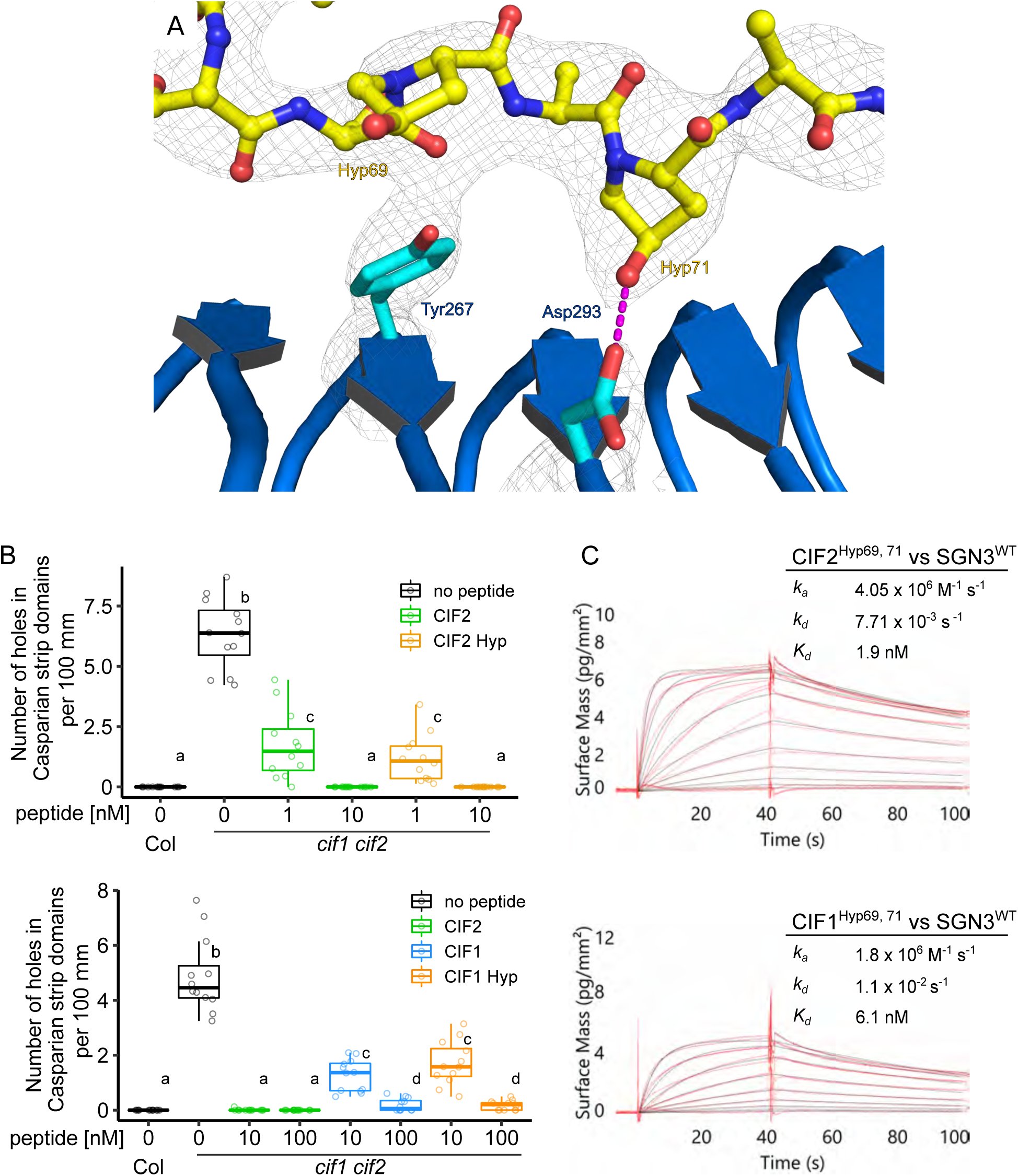
Two hydroxylprolines in CIF2 play no major roles in GSO1/SGN3 binding. (*A*) Details of the interaction between hydroxyproline residues of CF2^Hyp69, 71^ (yellow, in bonds representation) and the SGN3 ectodomain (blue ribbon diagram). Hydrogen bonds are depicted as dotted lines (in magenta), a 2F_o_-F_c_ omit electron density map contoured at 1.5 *σ* is shown alongside (gray mesh). (*B*) Quantitative analyses of number of holes in Casparian strip domains per 100 μm in *cifl cif2* double mutants treated with CIF peptide-variants (n=12 for each condition). Different letters indicate statistically significant differences (p <0.05, one-way ANOVA and Tukey test) (*C*) GCI assays of hydroxyprolinated CIF variants versus SGN3 wild type ectodomain. Sensorgrams are shown with raw data in red and their respective fits in black. Table summaries of kinetic parameters are shown alongside (k_a_, association rate constant; k_d_, dissociation rate constant; K_*d*_, dissociation constant).

**Fig. S7.**
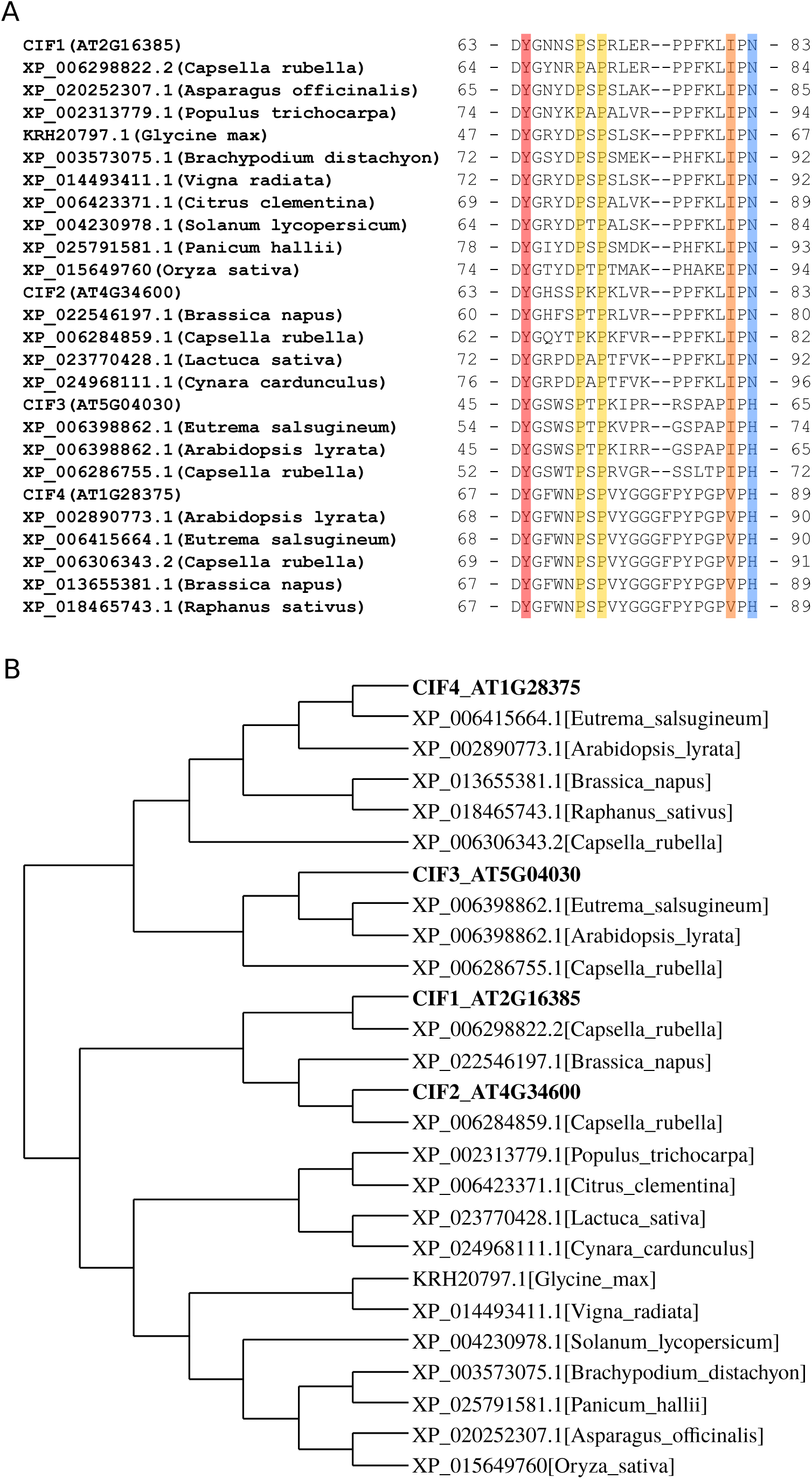
CIF3 and CIF4 orthologs are present in other plant species. (*A*) Multiple sequence alignment of CIF1-4 from Arabidopsis thaliana and their putative orthologs from other plant species. Sequences were obtained from NCBI (https://www.ncbi.nlm.nih.gov/) and aligned with the program T-coffee (version 12.0) (51). The conserved sulfated tyrosine is highlighted in red, hydroxyprolines in yellow, the conserved isoleucine in orange, and the C-terminal asparagine or histidine residue in blue. (A) Phylogenetic tree of CIF peptides prepared with the program BIONJ (52).

**Fig. S8.**
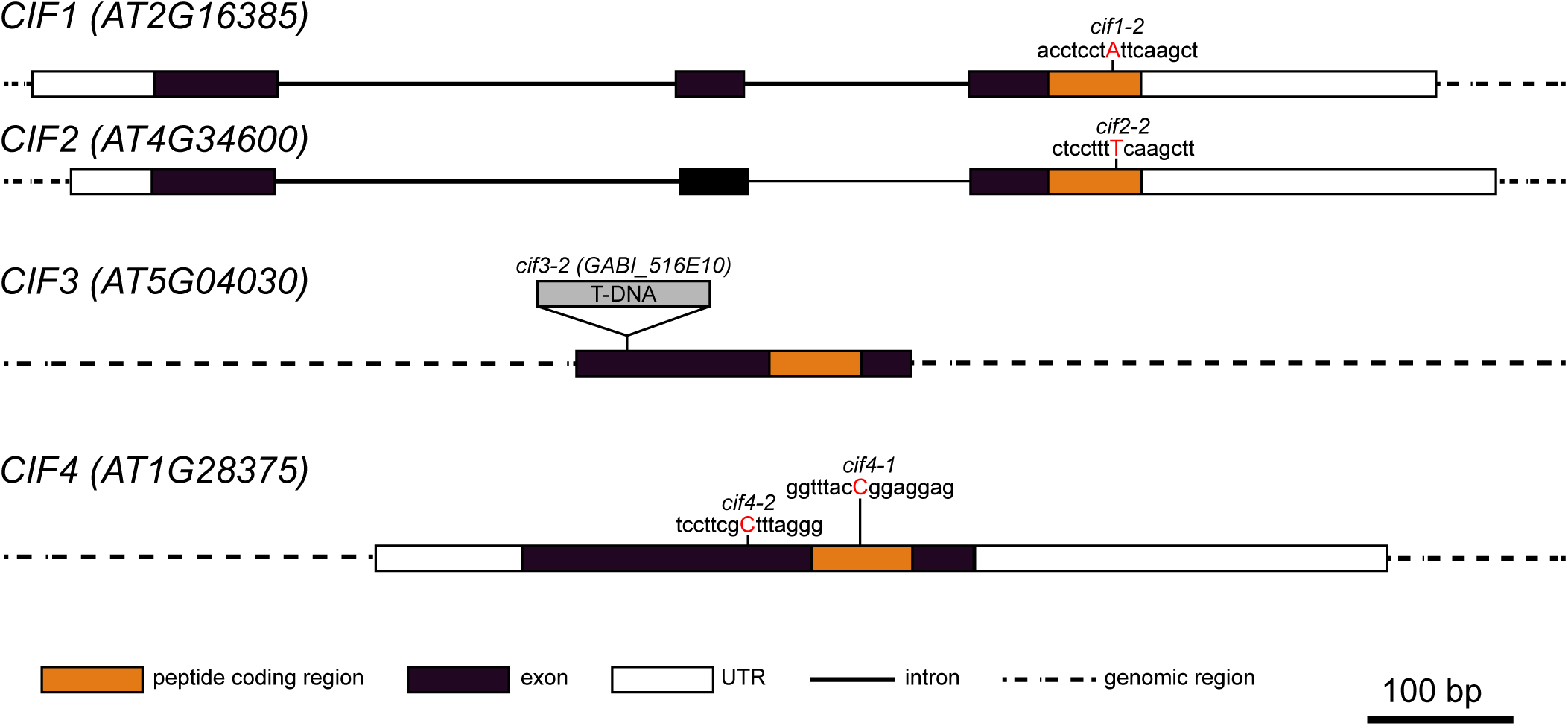
Overview of the CIF mutant alleles used in this study. Schematic models of the CIF genes and their mutant alleles. Single base pair insertion points (indicated by red uppercase letters) are shown together with their neighboring sequences. The T-DNA (gray box) insertion point is indicated in CIF3 locus.

**Fig. S9.**
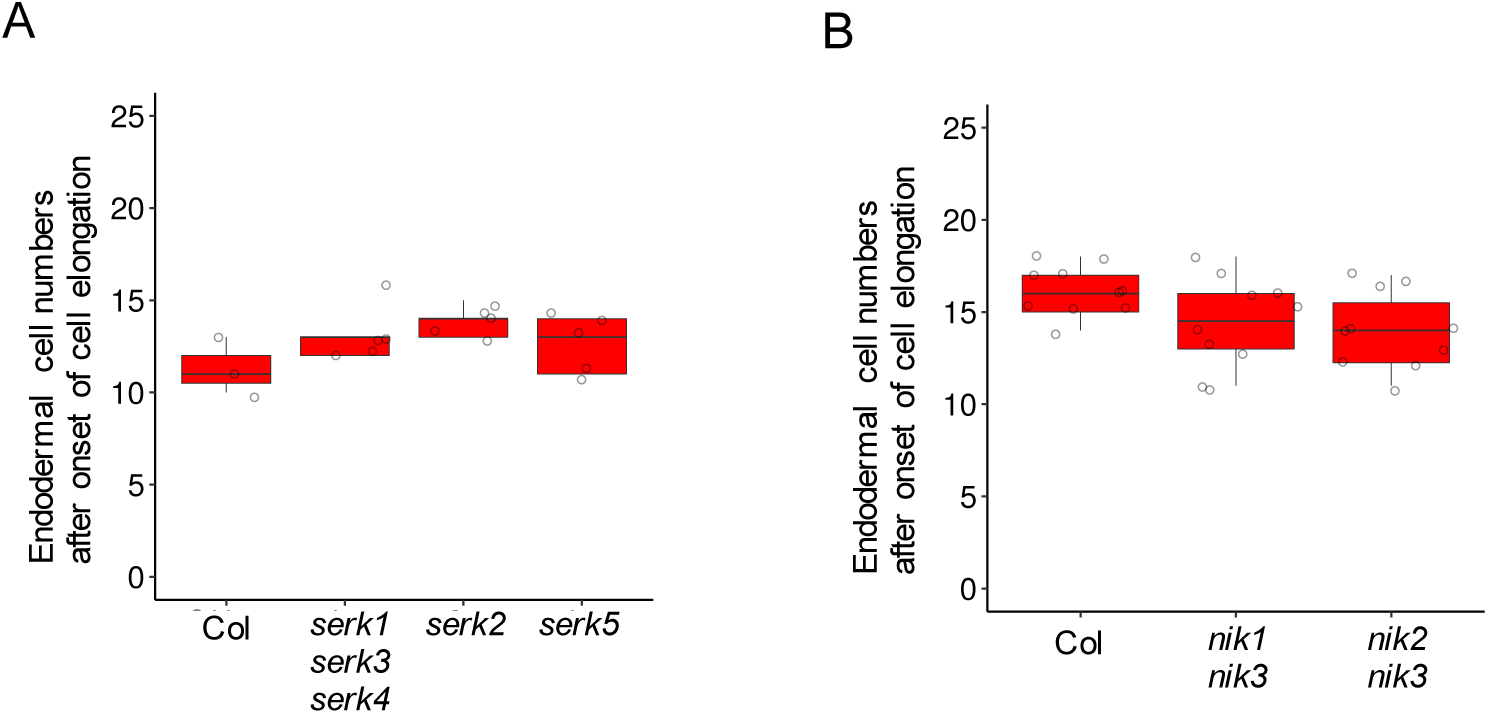
A number of *serk* and *nik* co-receptor loss-of-function mutants display no apparent Casparian strip defects. PI penetration assay with several *serk* and *nik single andIor multiple* mutants. Barrier functions were scored by counting the cell numbers until PI became impermeable to the steles.

**Fig. S10.**
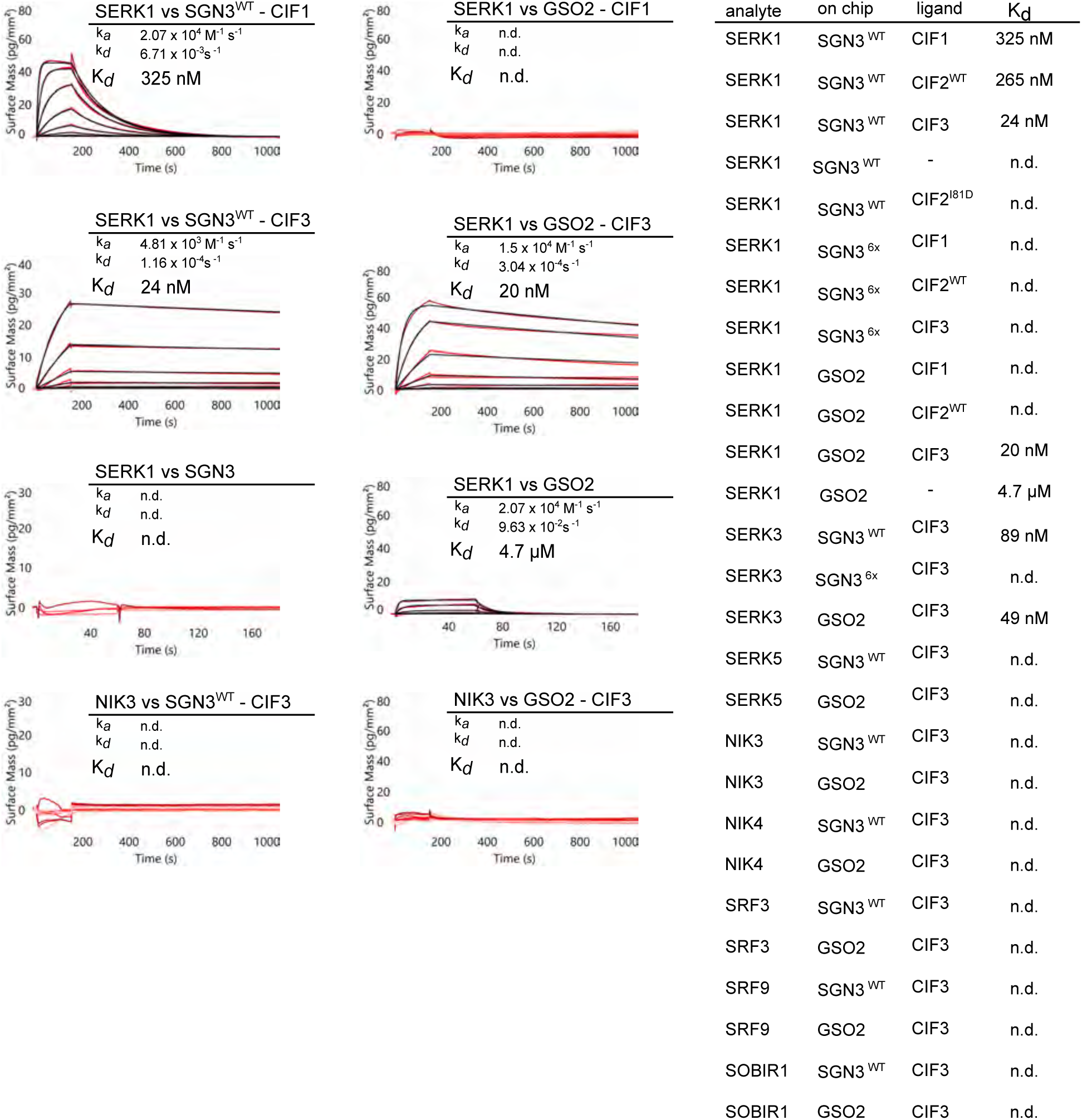
GSO1/SGN3 and GSO2 bind SERK1 and 3 co-receptor kinases in the presence of CIF peptides. GCI assays of co-receptor candidates versus GSO1/SGN3 and GSO2 ectodomains in the presence of CIF peptides. Sensorgrams are shown with raw data in red and their respective fits in black. Table summaries of kinetic parameters are shown (k_a_, association rate constant; k_d_, dissociation rate constant; K_*d*_, dissociation constant; n.d., no detectable binding).

**Fig. S11.**
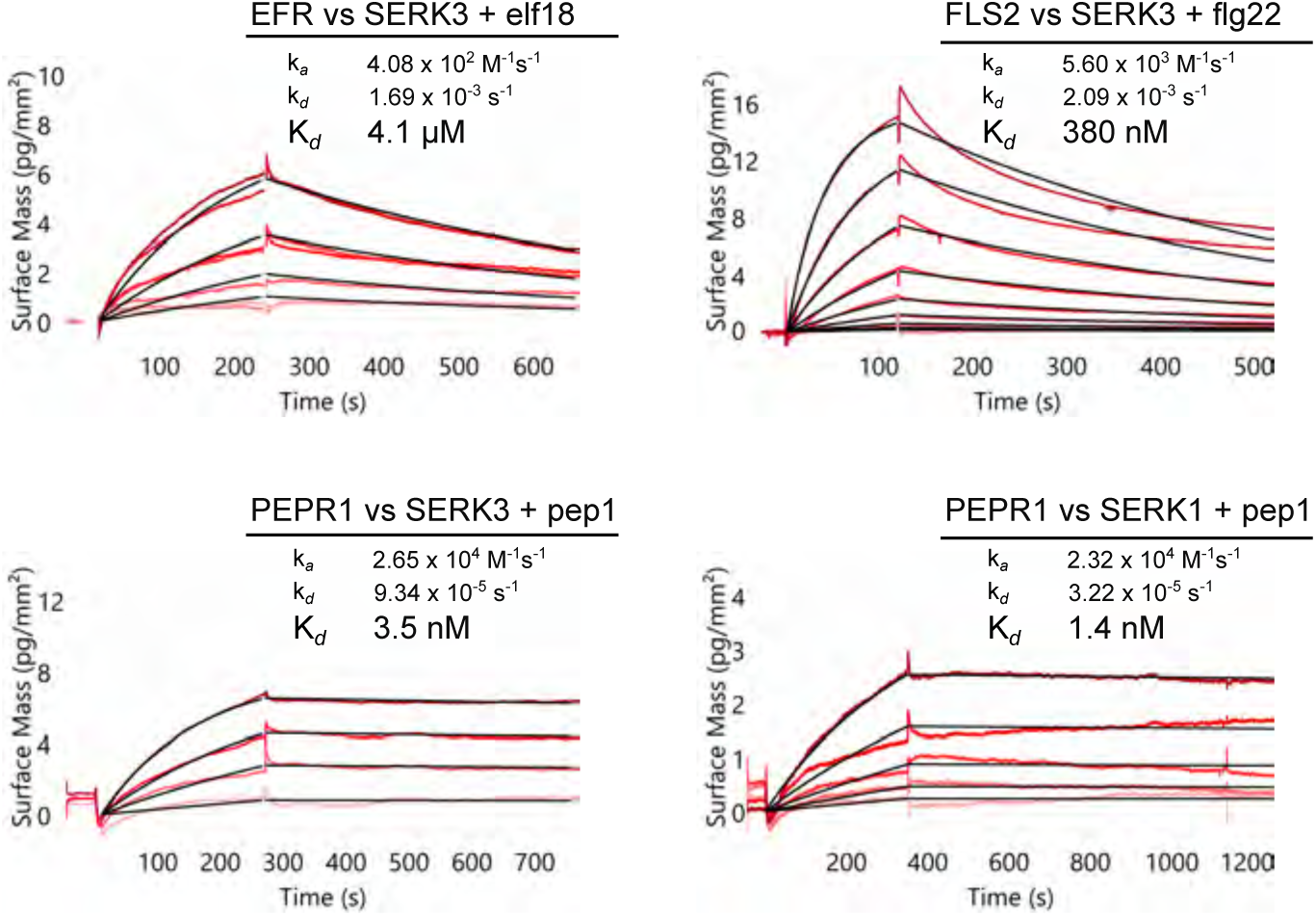
The LRR-RKs EFR, FLS2, PEPR1 bind SERKs with very different binding affinities and -kinetics. GCI assays of SERK co-receptors versus different, known LRR-RKs in the presence of their cognate peptide ligands. Sensorgrams are shown with raw data in red and their respective fits in black. Table summaries of kinetic parameters are shown (k_a_, association rate constant; k_d_, dissociation rate constant; K_*d*_, dissociation constant).

**Fig. S12.**
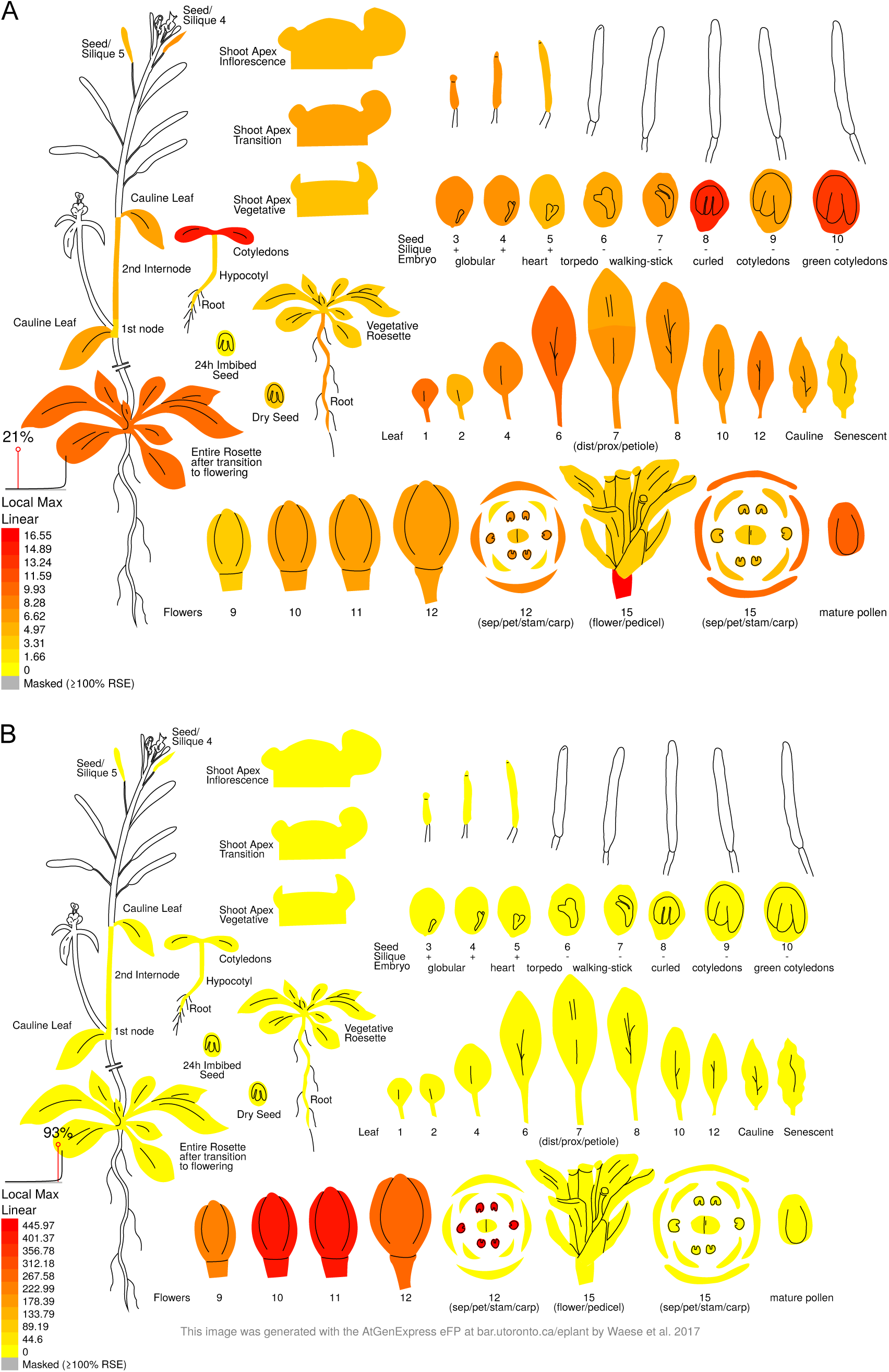
Expression analysis suggests putative functions for CIF3 and CIF4 outside Casparian strip formation / embryo development. Expression-pattern images of CIF3 (*A*) and CIF4 (*B*) were generated with the AtGenExpress eFP (https://bar.utoronto.ca/eplant/, (53)) using the publically available microarray data (54, 55). CIF3 appears to be expressed at embryo stage and in cotyledons, while CIF4 shows strong expression in early stage flowers and in stamens.

**Table S1.**
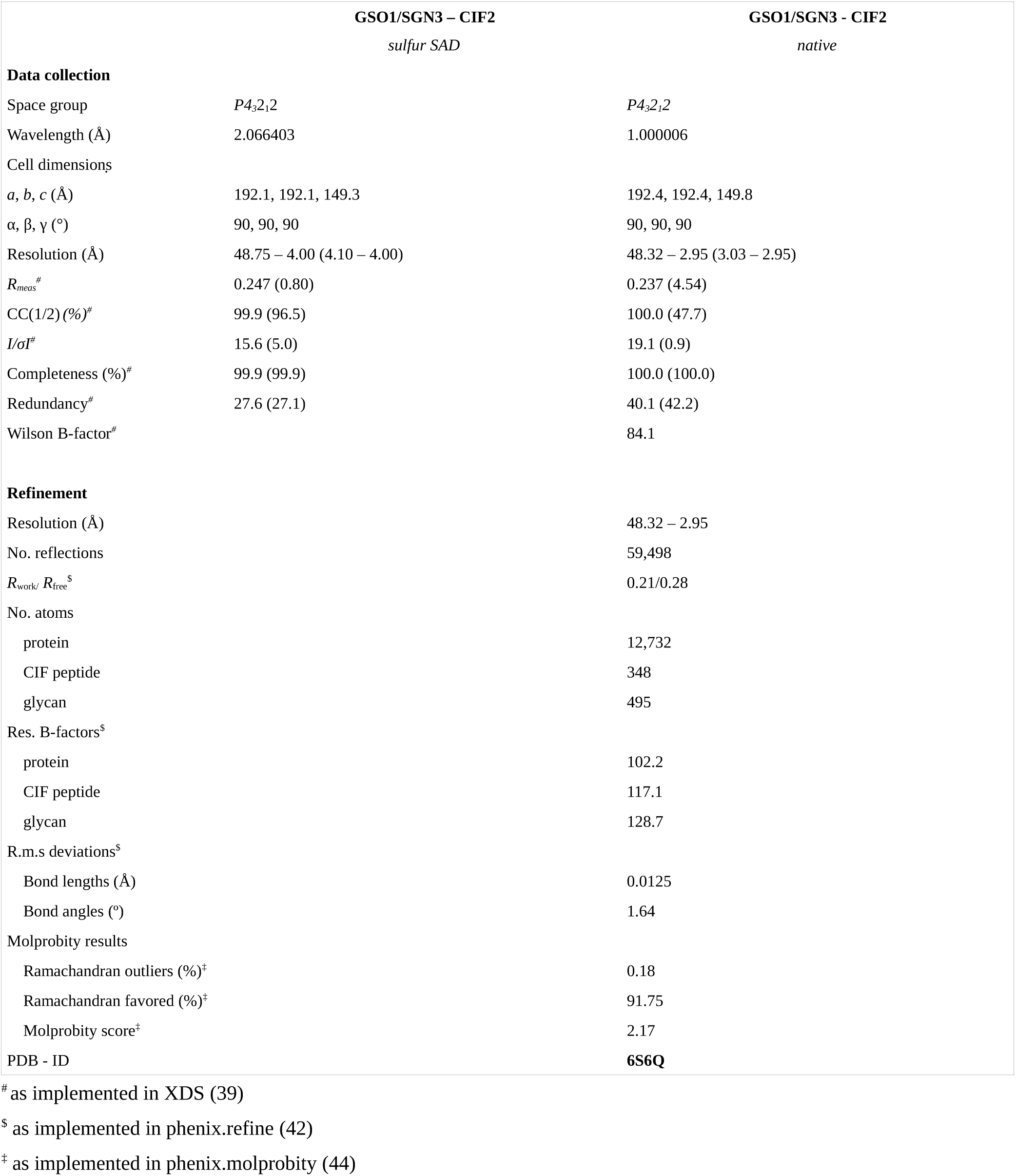
Crystallographic data collection and refinement

## Notes

#### Summary of Updates

Discussion updated, figure conversion errors fixed.

